# Models show altered outcomes from evolution of standing genetic variation in response to a suppression gene drive

**DOI:** 10.64898/2026.11.23.768563

**Authors:** Xinyue Zhang, Zichao Ma, Xiaohan Xie, Moming Guo, Jackson Champer

**Affiliations:** Center for Bioinformatics, School of Life Sciences, Center for Life Sciences, Peking University, Beijing, China; Department of Biosystems Science and Engineering, ETH Zürich, Basel, Switzerland

**Keywords:** Suppression gene drive, mosquito, genetic diversity, evolution, population genetics, computational simulation

## Abstract

Gene drives are commonly modelled as engineered inheritance-biasing systems with fixed performance, showing great potential for controlling disease vectors and other pests. Yet, natural populations contain standing genetic variation potentially capable of modifying drive transmission, resistance allele formation, and drive-associated fitness. This could lead to evolution in the population after the release of a drive system, potentially affecting the outcome. Here we use individual-based panmictic and spatial mosquito models to examine how standing genetic variation responds to the release of a homing drive configured for population suppression. Genetic variation affecting drive conversion efficiency showed a robust anti-drive response, evolving towards reduced conversion and resulting in inferior outcomes if rapid population elimination was avoided. In contrast, variation affecting embryo resistance and drive fitness cost showed context-dependent evolutionary responses associated with the spatial dynamics of the drive. These could evolve to either increase and inhibit the drive or decrease and improve drive outcomes. Evolving genetic variation in dispersal slightly increased successful elimination outcomes. Recessive deleterious variation acting through inbreeding depression reduced the minimum post-release population size, potentially increasing population elimination rates. However, its effects were not effective in the long-term since deleterious mutations were quickly purged among populations that persisted. Together, these results identify component and scenario-specific evolutionary trajectories based on drive pressure and available standing genetic variation. Thus, standing genetic variation may be treated as a dynamic part of theoretical gene drive modeling and could be worth experimentally quantifying for some possible release candidates.

## Introduction

Mosquitoes transmit a variety of pathogens, including malaria, dengue, yellow fever virus, zika virus, etc., causing serious public health issues worldwide (WHO, 2026). Vector control still relies largely on chemical insecticides, but this approach is increasingly constrained by insecticide resistance and concerns about environmental pollution. In recent decades, gene drive systems have been considered as a potential strategy for controlling disease vectors and other pests (Bier, 2021; Chen et al., 2026; Hay et al., 2021; Verkuijl et al., 2022; Zhang et al., 2026). Gene drives can bias their own inheritance, enabling them to spread rapidly through populations (Bier, 2021; DiCarlo et al., 2015; Harvey-Samuel et al., 2023). They can be used for population modification with cargo genes that prevent disease transmission (Adelman and Kojin, 2021; Carballar-Lejarazú and James, 2017; Fuchs et al., 2013; Reid et al., 2021) or directly suppress vector populations (Champer et al., 2021; Champer et al., 2022; Kyrou et al., 2018). However, technical and regulatory hurdles constrain field experiments (Kc et al., 2020; Kormos et al., 2022), making computational models important for anticipating their performance in natural environments.

Suppression drives are generally more sensitive to performance parameters compared to modification systems. They generate population density variation across a continuous landscape, creating conditions for chasing dynamics and altering outcomes relative to panmictic models (Champer et al., 2021; Girardin and Débarre, 2021; North et al., 2019; Paril and Phillips, 2022). When the drive eliminates a population locally, adjacent wild-type individuals can recolonize the resulting low-competition space, forcing the drive to “chase” wild-type individuals across the landscape (Birand et al., 2022; Champer et al., 2021). Chasing dynamics can cause periodic variation in population density across time and space and may lead to population elimination, drive loss, or long-term unstable coexistence. This can be a major cause of drive failure for otherwise efficient systems in spatial models (Bull et al., 2019; Champer et al., 2021; Olejarz and Nowak, 2024; Zhang et al., 2024). These long-term dynamics provide an opportunity for functional resistance allele formation, even if it could be otherwise avoided. They could also allow more time for selection based on standing genetic variation. The resulting evolutionary response may in turn alter drive performance.

In genetically diverse strains, the same drive system can show substantial variation in drive conversion and embryo resistance allele formation. Across 128 strains of the *Drosophila* Genetic Reference Panel (DGRP), embryo resistance allele formation varied by more than an order of magnitude (Champer et al., 2019). That study identified several genes that affect embryo resistance allele formation using GWAS and short-hairpin RNA knockdown, but it also suggested that drive-related traits may involve multiple loci of relatively small effect rather than a few large-effect genes. However, such effects were not consistently detected across species and drive systems. In *Anopheles gambiae*, homing was maintained across the genetic backgrounds and chromosomal inversion arrangements examined (Pescod et al., 2023), perhaps because it was very high in the first place. A study of European *Drosophila melanogaster* strains likewise observed variable homing efficiency for an intermediate-efficiency system but detected no statistically significant heritable genetic component (Faber et al., 2025). Potential molecular contributors to background-dependent variation include differences in Cas9 expression and persistence, as well as DNA repair processes (Champer et al., 2017; Champer et al., 2019). The extent to which these processes explain the observed differences in drive performance remains unresolved. Studies of quantitative traits indicate that genes outside canonical pathways can influence pathway function and that many small-effect variants can cumulatively alter trait performance (Hua and Springer, 2018).

Gene drive theory has largely emphasized standing and newly formed resistance at the target site (Khatri and Burt, 2022; Unckless et al., 2017), but standing variation elsewhere in the genome may make other components of drive performance evolutionarily labile. A mutation affecting drive conversion, embryo resistance formation, or drive fitness could be selected for or against. We therefore expect component-specific evolutionary responses. Some components may evolve in a stable direction across regimes, whereas others may reverse as drive pressure and genotype composition change.

There is another type of genetic diversity pre-existing in natural populations that could influence suppression drive outcomes and population dynamics. Inbreeding increases homozygosity, which can expose recessive deleterious alleles that were masked in heterozygotes. The resulting decline in survival or reproduction is one genetic component of inbreeding depression (Charlesworth and Willis, 2009). A suppression drive may intensify this process by reducing local or population-wide abundance, thereby increasing related matings and exposing previously hidden load precisely when demographic recovery is most important. Previous modelling has shown that inbreeding can impede drive spread by increasing related matings or altering genotype production (Bull, 2016; Bull and Barrick, 2017; Bull et al., 2019; Champer et al., 2021). Its ultimate effect may nevertheless depend on the severity of inbreeding depression: weak depression can favor and even evolve to inbreeding strategies that obstruct a drive (Bull, 2016), whereas stronger depression can deepen population decline and increase successful elimination in some settings (Champer et al., 2021). These results motivate a mechanistic treatment of the genetic load underlying inbreeding depression rather than a fixed mating or fitness penalty. Previous gene drive models have only represented inbreeding or inbreeding depression through a fixed fitness penalty or mating parameter (Bull et al., 2019; Champer et al., 2021). However, standing recessive load is dynamically exposed and purged (Grossen et al., 2020; Hedrick and Garcia-Dorado, 2016), which could increase the chance of population recovery.

Because gene drive creates a novel selective environment that can recruit preexisting genetic variation, we hypothesized that standing mutations affecting drive conversion, embryo resistance allele formation, fitness costs, and dispersal can show directional evolutionary responses and can change the outcome of a suppression drive release. We tested how genetic architecture influences these mutation responses. Separately, by explicitly simulating genome-wide recessive deleterious variants before and after release, we examined whether drive-induced decline exposes hidden load, deepens the initial population bottleneck, and possibly increases the chance of successful population elimination before purging the genetic variation. By modelling standing genetic variation, rather than *de novo* mutations that accumulate after release, we aim to make predictions of gene drive dynamics more relevant to natural populations, showing what changes might be possible and when they might occur. These interactions identify the drive properties and population contexts that could be considered when evaluating potential gene drive release candidates.

## Methods

### Mosquito model

Models were implemented in the SLiM forward-time evolutionary modelling framework (version 4.1) (Haller and Messer, 2023). Simulation outputs were processed and analyzed in Python 3.10.

In this study, we use an individual-based population model simulating *Anopheles* and other mosquitoes based on our previous study (Champer et al., 2022). The life cycle of mosquitoes is modeled as weekly time-steps. Female mosquitoes have longer lifespans than males. We set the maximum lifespan of female mosquitoes as 8 weeks, and males as 5 weeks, and the first two weeks are juvenile stages (egg, larvae, and pupa). The number of adult individuals of each age linearly decreases to zero through time, which is consistent with survival as tested in the laboratory (Christiansen-Jucht et al., 2014). Adult females search for mates if needed and have a 50% probability to reproduce in every time step. Mated females store sperm for reproduction in future weeks, but they also have a 5% chance to remate and update the sperm storage in each week (Gomulski, 1990). The number of offspring produced in a week where reproduction occurs is drawn from a Poisson distribution with an average *λ*=2∗*β_max_*∗*ω_i_*, where *β_max_*represents the maximum number of female offspring per female, and *ω_i_* represents the genotype fitness of the female that is reproducing. After reproduction, age 0 larvae compete for food supply and other development resources to survive while adults do not need to compete (Ross et al., 2014; Walsh et al., 2011). Thus, population density is regulated by larval survival rather than adult fecundity. Because larger larvae consume more resources than smaller larvae, week-old larvae contribute five times as much competition as new larvae. We simplify their survival as 100% (so larvae are eliminated at age 0 unless they are destined to develop into adults). The survival rate (*v*) of an age 0 larva is dependent on the ratio of actual number of larvae to expected number of larvae at capacity, which is expressed as:

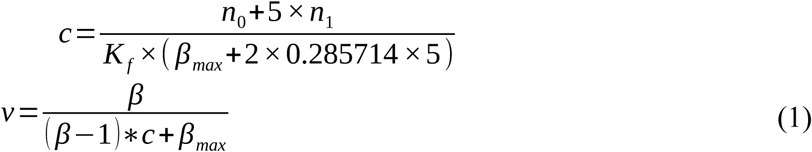

Where *n_0_* and *n_1_* represent the number of larvae of age 0 and 1, respectively, *K_f_* represents the number of adult females when the population size is at capacity and the population is at equilibrium, 0.285714 indicates the proportion of age 1 larvae when the population is at equilibrium, and *β* represents the highest growth rate of the population when at low density (normally set to 6).

### Spatial model

The spatial model simulates a two-dimensional continuous landscape in which each individual’s position is recorded. Wild-type individuals were initially distributed at random in a square with side length 1, corresponding to 3.4 km. Carrying capacity was set to K = 16,000 total individuals. This reduced size was used for computational tractability and should not be interpreted as the absolute field census implied by the reported density of 430 adults per hectare (Yao et al., 2022). However, actual effective population sizes are substantially lower than actual sizes, so this reduced level may better represent real-world population dynamics than a model using the full census size without additional compensation. Each week, unmated or remating females select a mate at random within a radius equal to the mean dispersal distance. This distance was set to 0.03 landscape units, corresponding to approximately 101 meters per week (Yao et al., 2022). If females cannot find a mate, they will not reproduce.

The spatial model used a resource-explicit model in which competition between individuals is determined by the number of abstract resources and the number of individuals sharing these resources (Champer et al., 2024), which gives a better runtime performance and nearly identical results compared to models that directly simulate interactions between individuals. Resource nodes are uniformly hexagonally tiled across the landscape. Each resource node can serve larvae that are less than the foraging distance away from it, and there are on average four resource nodes in an individual’s foraging area. Resource nodes discretize local competition rather than directly specifying carrying capacity. Demand from one individual is spread between all nodes in range. An average of four nodes per foraging area was used to balance spatial resolution and computational cost. Population abundance remained stable in no-drive control simulations under this setting. The foraging area of larvae is much less than the dispersal area of adults due to motility differences, so we set a separate foraging distance of 0.01 (actual larvae travel even less, but only a small fraction of the habitat is suitable for larvae). For each age-0 larva, we found the average weighted larval counts across all resource nodes within its foraging range. At each node, age-0 and age-1 larvae contributed weights of 1 and 5, respectively. This mean resource demand was normalized by the expected competition level to obtain the relative competition intensity *c*, producing the same competition as our panmictic model:

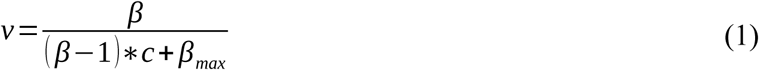

Surviving adult mosquitoes migrate with a random direction and a distance drawn from the distance kernel, which is the probability density function:

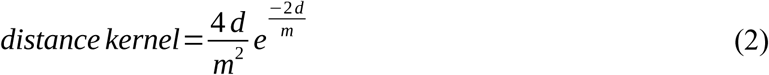

where *d* represents the average dispersal distance. This negative exponential distance kernel has been supported by mark-release-recapture experiments and spatial genetics tests (Filipović et al., 2020; Winskill et al., 2015). The positions of offspring are offset from their mother with a distance drawn from the same distance kernel and a random bearing. If any individual is placed outside the arena, its position is redrawn until it falls within the arena. Nearest neighbor index (NNI) is used to assess spatial population dynamics (Zhang et al., 2024).

### Suppression strategy

Female fertility homing drives can sterilize females to suppress a population, and they have already been implemented in laboratory settings (Kyrou et al., 2018; Yang et al., 2022). CRISPR/Cas9 targets a haplosufficient but essential female fertility gene, disabling it with the drive itself or with nonfunctional resistance alleles. After cleavage happens in the germline of a drive/wild-type heterozygote, the wild-type allele can be converted to a drive allele by homology-directed repair with a probability equal to the drive conversion rate. It could also be converted into a germline resistance allele by end-joining repair with a chance equal to the germline resistance allele formation rate. Resistance alleles refer to alleles that cannot be cleaved by Cas9 guided by the designed gRNAs. Maternal deposition of Cas9 and gRNA could additionally convert wild-type alleles in new embryos (regardless of which parent they are received from) to resistance alleles at a rate equal to the embryo resistance allele formation rate.

There are two types of resistance alleles, “r1” and “r2”, corresponding to whether or not the target gene function is maintained. Functional “r1” resistance alleles play a decisive role in preventing population suppression. Even if they form at a low probability, the population is likely to recover, resulting in drive elimination (Birand et al., 2022). In our simulations, we assume that end-joining repair only results in nonfunctional alleles by using multiplexed gRNAs (Champer et al., 2020; Morianou et al., 2026; Yang et al., 2022) and conserved target sites (Kyrou et al., 2018). Females were sterile when neither target gene copy retained wild-type function. In addition, unintended somatic expression of Cas9 or disruption of the target gene in germline cells could also reduce reproduction in female drive heterozygotes. We represented this cost by the female drive heterozygote fitness parameter, with wild-type relative fitness set to 1.

The initial drive release number does not influence the final suppression level and outcome in most situations because the drive has no introduction threshold and increases rapidly. We therefore released 0.1K male drive heterozygotes (K = 16,000 total individuals); in spatial simulations, all released males were placed within a central circle of radius 0.01. Simulations continued for 1,500 weeks (approximately 474 generations) unless population elimination or drive loss occurred earlier. Long-term persistence outcomes were recorded when drive and non-drive alleles coexisted at week 1,500.

### Genetic variation

Genetic elements were allocated between chromosomes 2 and 3 in proportion to their annotated gene counts in Ensembl Metazoa release 60 for the AgamP4 assembly, and the total number is set as 10,000 (Melo and Wallau, 2020; Yan et al., 2012). The genetic lengths of the second and third chromosome were set as 72.1 cM and 93.7 cM, referring to the genetic map of *Anopheles gambiae* (Zheng et al., 1996), and the chromosome-specific genetic map was divided across adjacent element intervals to obtain the recombination rate per interval. The drive allele was placed at the center of the second chromosome. Mutations that influence different characteristics were randomly distributed on these loci.

Standing variants affected drive conversion, embryo resistance formation, or female drive heterozygote fitness through additive effects. Each trait was examined separately against controls carrying neutral variants. We also varied the number and effect sizes of modifier loci to distinguish the contribution of realized standing variation from genetic architecture. Genomic settings and variant initialization are described in Supplemental Methods S1.

For dispersal, we introduced heritable variation and a small fitness cost of greater movement. Evolving-dispersal populations were compared with matched fixed-dispersal controls and no-drive references (Supplemental Methods S2). To examine recessive deleterious variation, we allowed fully recessive mutations to affect juvenile viability and established standing variation before drive release. We followed mutation burden and assessed potential inbreeding depression using virtual matings that did not contribute offspring to the simulated population (Supplemental Methods S3). Default parameters were partially based on previous studies of gene drive and other genomic variation.

### Data generation

Code and source data are available on GitHub (https://github.com/xinga1516/evolution-to-gene-drive). For most simulations, variable mutation and neutral mutation control simulations were generated independently, without matched random seeds and under the same mutation generation and genetic settings. Mutation alleles altered the focal drive trait only in the functional treatment. Their effect value was recorded but did not alter the focal drive trait in the control. For deleterious load and variable dispersal simulations, otherwise corresponding simulations were run with and without variations. Weekly outputs included allele frequencies, total population size, nearest-neighbor index for spatial simulations, population mean mutation effect, and the measurements required for outcome, minimum-population-size, and deleterious allele purging.

### Statistical analysis

Independent simulations were the units of analysis. We compared independent continuous endpoints with permutation tests, selected unpaired outcome contrasts with Fisher’s exact tests, and matched dispersal outcomes with exact McNemar tests. Dispersal changes were compared with matched no-drive references using paired t-tests. Setting-level associations used Spearman’s rank correlation. Genetic architecture analyses used linear interaction models with HC3 robust inference and joint Wald tests. Continuous-endpoint intervals were estimated by bootstrap, and outcome probabilities used Wilson intervals. Holm correction was applied within specified test families. Tests were two-sided where applicable, with significance assessed at *P* < 0.05. Full procedures are given in Supplemental Methods S4.

## Results

### General framework and variable drive parameter settings

When analyzing the effect of standing genetic variation, we chose four simulation settings. Two of these were in panmictic populations, and two were in spatial populations, each with high and low drive performance. These treatments generated distinct distributions of population elimination, drive loss (a rare outcome), and long-term persistence. The less efficient drive would usually result in an equilibrium outcome in both panmictic and spatial settings. The higher performance system would usually result in panmictic population elimination and “chasing” in the spatial population (recorded as long-term persistence). By having each of these qualitatively different situations, we could assess how standing genetic variation would evolve in each setting. We used control simulations with segregating neutral mutation loci that do not alter the focal trait.

### Variable conversion mutations show a directionally robust anti-drive response

We first defined four experimental regimes by crossing population structure (panmictic or spatial) with drive conversion (0.70 or 0.85, with other parameters at their default values in Table S1). The higher genetic load associated with the 0.85 drive conversion efficiency regime was sufficient to eliminate the panmictic population, whereas the lower genetic load under the 0.70 regime was insufficient, resulting predominantly in long-term persistence (**Figure 1A**). In spatial populations, both drive regimes were instead dominated by long-term persistence. When standing genetic variation affected drive conversion, ultimate outcomes were not altered, but significant evolution did take place. Variable mutations produced negative responses under all four settings, though it had only a minor effect in the high performance panmictic scenario when population elimination took place quickly. In others, average drive conversion efficiency was significantly reduced toward the end of the simulations, falling approximately 20% (two-sided permutation tests, all adjusted *P* <0.001; **Figure 1B**, **Table S2**). When sufficient post-release exposure was available, variable conversion mutations thus evolved consistently towards reduced transmission. For all simulations with long-term persistence, the evolution of drive efficiency had the practical effect of substantially increasing the equilibrium population (**Figure 1C**). The reduced genetic load of the drive was particularly apparent in the panmictic population, but the spatial populations also were able to rebound substantially. Among spatial simulations ending in long-term persistence, the low performance drive showed comparatively stable NNI trajectories, whereas the 0.85 drive conversion simulations with neutral mutations were different than those with variable mutations (**Figure 1D**). Specifically, the NNI trajectory shifted from with chasing character to a higher and more stable value more closely associated with an even equilibrium when drive conversion was 0.85 in the spatial model (**Figure 1D**). The mean NNI of final 20 weeks also showed significantly different values between neutral control and groups with variable mutations (**Figure S1**).

**Figure 1.**
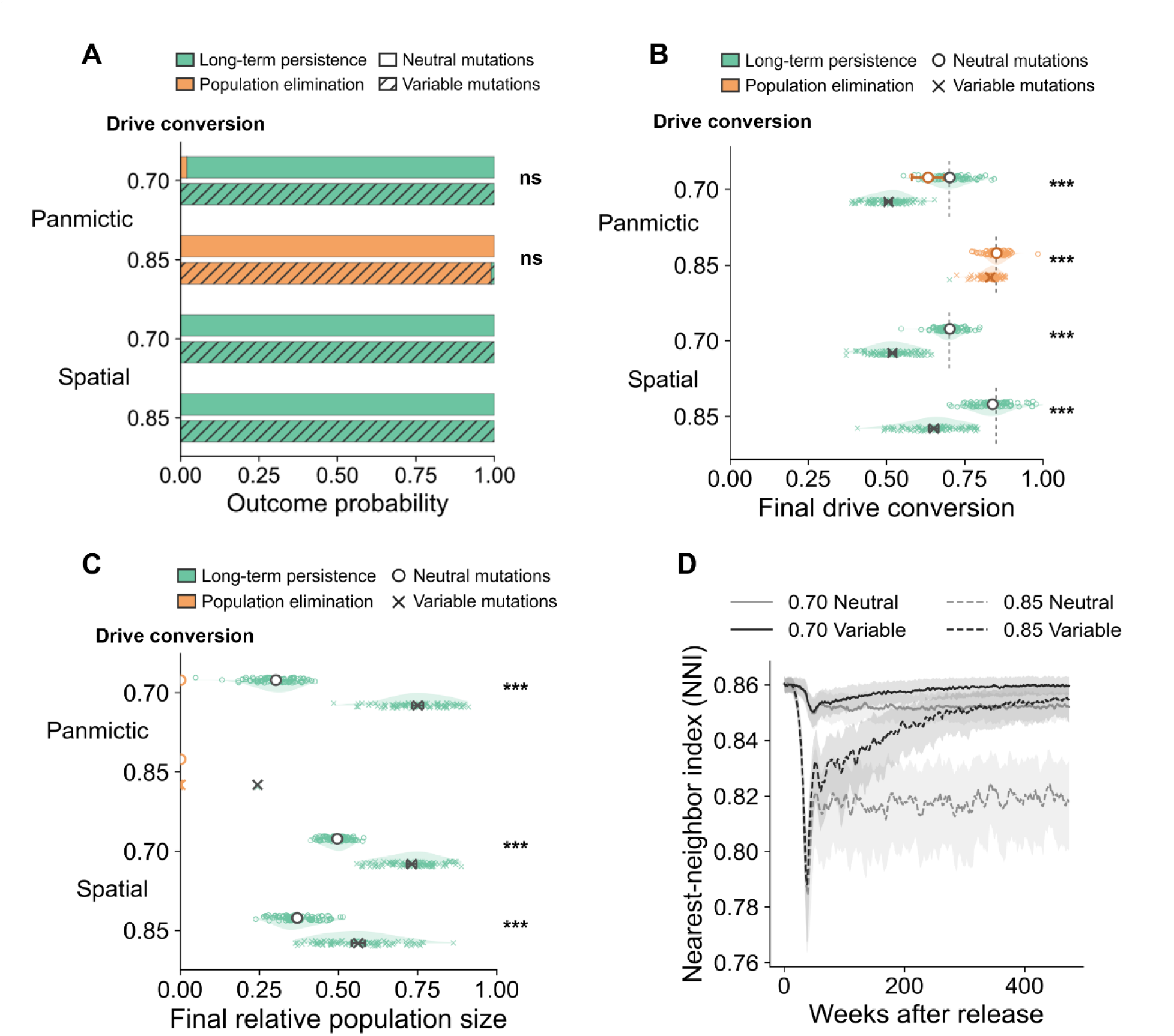
Effect of standing genetic variation influencing drive conversion efficiency. (**A**) Outcome composition across all simulations. Each group contained 100 simulations. Wilson 95% confidence intervals for outcome probabilities are reported in the Source Data. (**B**) Final mean drive conversion of last 20 weeks in independent variable-mutation and neutral-mutation control simulations. Half-violins and small points show the replicate-level distributions. Large dots indicate means, and horizontal error bars show 95% confidence interval. Some horizontal error bars are too short to seen. Short dashed lines indicate the nominal initial values of 0.70 or 0.85. Exact n, mean differences and raw and adjusted *P* values are reported in Table S2. (**C**) Average relative long-term population size of the last 20 weeks compared to initial level among long-term-persistence simulations. Plotting conventions and analysis are as above. (**D**) Spatial NNI trajectories among long-term-persistence simulations; lines show medians, and ribbons show the 25th– 75th percentile interval. ns *P* ≥ 0.05; \**P* < 0.05; \*\**P* < 0.01; \*\*\**P* < 0.001.

### Embryo resistance mutation evolution reverses direction across drive regimes

For the analysis of embryo resistance mutation evolution, we used the same four parameter sets as the drive conversion analysis. In the panmictic simulations with a drive conversion efficiency of 0.7, variable embryo resistance mutations increased successful elimination from 3/100 to 15/100 simulations (two-sided Fisher exact test, unadjusted *P* = 0.0208; **Figure 2A**). Relative to the baseline, embryo resistance mutations showed a qualitatively different evolutionary response across drive regimes. In most simulations (except those where the population was rapidly eliminated in panmictic populations with high drive efficiency), average population embryo resistance declined (two-sided permutation tests, all Holm-adjusted *P* < 0.001; **Figure 2B**). However, the direction of embryo resistance evolution reversed in the high drive efficiency (initially chasing) spatial regime. Note that evolutionary responses were more variable among individual simulations, particularly in the more dynamic spatial regimes, and a small number of persistent simulations changed in the opposite direction to their regime-level mean (**Figure 2B**).

**Figure 2.**
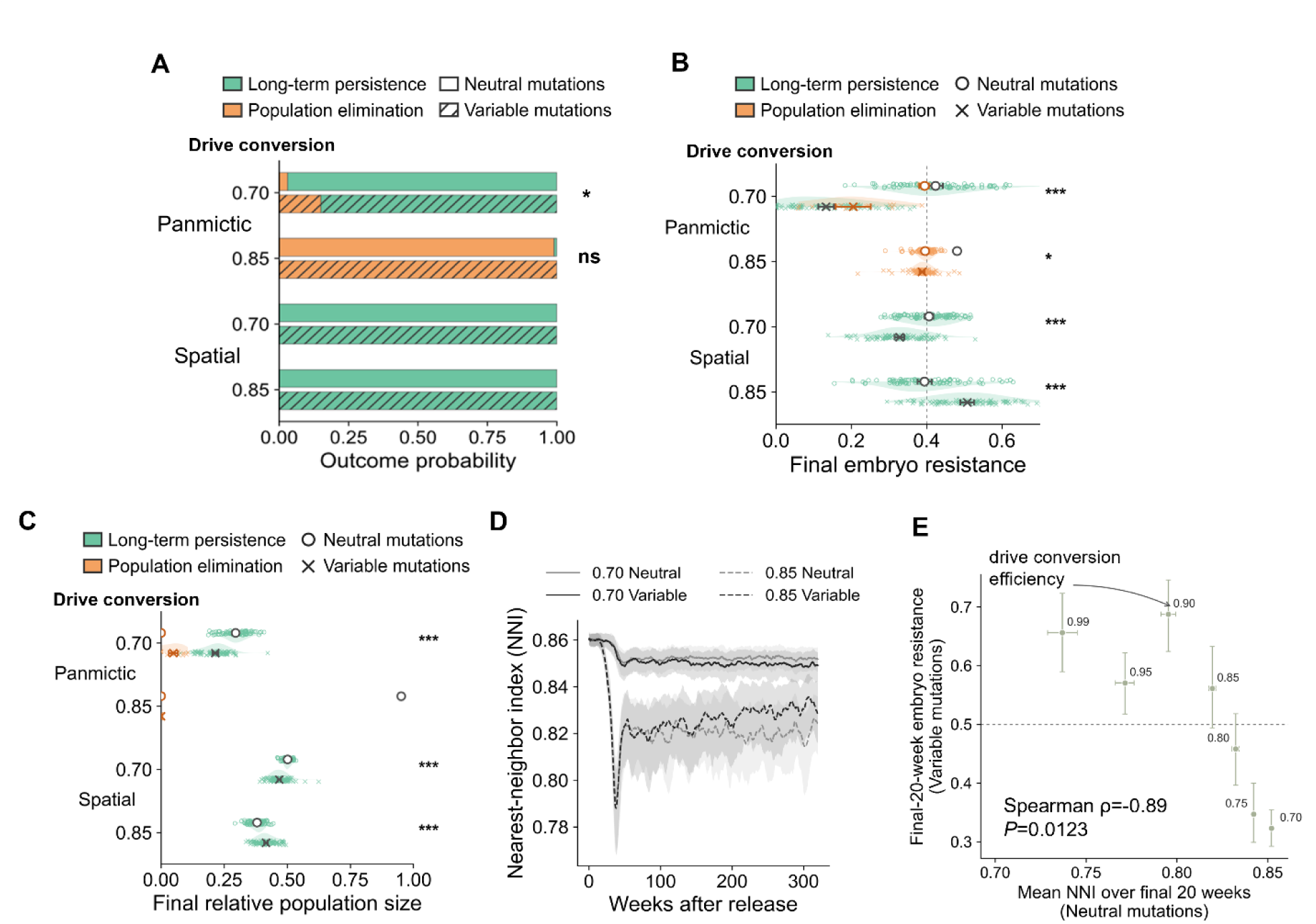
Effect of embryo resistance allele formation rate mutations on suppression gene drive. (**A**) Outcome composition across all simulations. Wilson 95% confidence intervals for outcome probabilities are reported in the Source Data. The Panmictic-0.70 elimination comparison was evaluated using a two-sided Fisher exact test (*P* = 0.00519). (**B**) Final embryo resistance allele formation rate in independent variable-mutation and neutral-mutation control simulations. The endpoint for each independent simulation was defined as the adult-population-size weighted mean of the final 20 population-level average-attribute records. Half-violins and small points show replicate-level distributions. Large dots indicate arithmetic means, and horizontal error bars show 95% confidence intervals. Some horizontal error bars are too short to seen. The continuous dashed line indicates the initial modifier attribute of 0.4. Variable and neutral groups were compared separately within each drive regime using two-sided independent two-sample permutation tests. Exact sample sizes, mean differences, and raw and adjusted *P* values are reported in Table S2. (**C**) Average relative long-term population size of the last 20 weeks compared to initial level among long-term-persistence simulations. Plotting conventions and bootstrap intervals are as above. (**D**) Spatial NNI trajectories among simulations ending in long-term persistence. Solid lines show medians and shaded ribbons indicate the 25th–75th percentile interval; (n=100) simulations were included in each group. ( **E**) Parameter-level association between the neutral-control mean NNI over the final 20 weeks and the corresponding mean final embryo resistance allele formation modifier attribute in variable mutation simulations across seven drive values. Each parameter setting contained 100 simulations. Error bars show bootstrap 95% confidence interval. The association was evaluated using a two-sided exact Spearman permutation test ( *ρ*=−0.892857, *P* = 0.0123). ns *P* ≥ 0.05; \**P* < 0.05; \*\**P* < 0.01; \*\*\**P* < 0.001.

Relative to neutral controls, variable-mutation simulations had significantly lower long-term population averages when drive efficiency was low, but significantly higher relative population size compared to neutral mutations for the high efficiency spatial model (two-sided permutation tests, all adjusted (*P* < 0.001); Figure 2**C**). The final population size was relatively reduced approximately 20% in the panmictic setting, but less so in the lower efficiency spatial model, while the high efficiency spatial model had a slightly increased population (Figure 2**C**). The persistent spatial treatments also followed different NNI trajectories for the high efficiency drive ( Figure 2**D**). Specifically, NNI increases, showing a reduction in chasing character toward a more even equilibrium. However, the final populations showed large variance for the higher efficiency drive, indicating that random factors have a large role in the population dynamics (**Figure S1B**).

To test whether embryo resistance evolution covaried with the spatial dynamics generated by different drive settings, we compared final embryo resistance with baseline across seven drive conversion efficiencies (0.70, 0.75, 0.80, 0.85, 0.90, 0.95, and 0.99 - values below 0.85 lack sufficient genetic load to eliminate panmictic populations). All simulations ended in long-term persistence, and neutral-mutation NNI was negatively correlated with the mean final embryo resistance attribute (Spearman’s *ρ*=¿-0.892857, *P* = 0.0123; Figure 2**E**). Embryo resistance was reduced only in simulations with lower NNI, indicating an even equilibrium state rather than chasing. Thus, embryo resistance evolution differed among experimentally defined regimes and was associated with their realized spatial dynamics.

### Fitness mutations show context-dependent responses

We next examined standing variation affecting the fitness of female drive heterozygotes across drive regimes. Here, we used a laboratory-informed high drive conversion parameterization with female and male efficiency at 0.99 and 0.96, respectively, germline resistance of 0.01 and 0.02, embryo resistance formation rate of 0.08, and nominal initial female heterozygote fitness of 0.5 (Champer et al., 2022; Kyrou et al., 2018). We also examined a lower efficiency drive with similar parameters but a drive conversion efficiency of 0.70, evaluating each in both panmictic and spatial models (Figure 3**A–D**).

**Figure 3.**
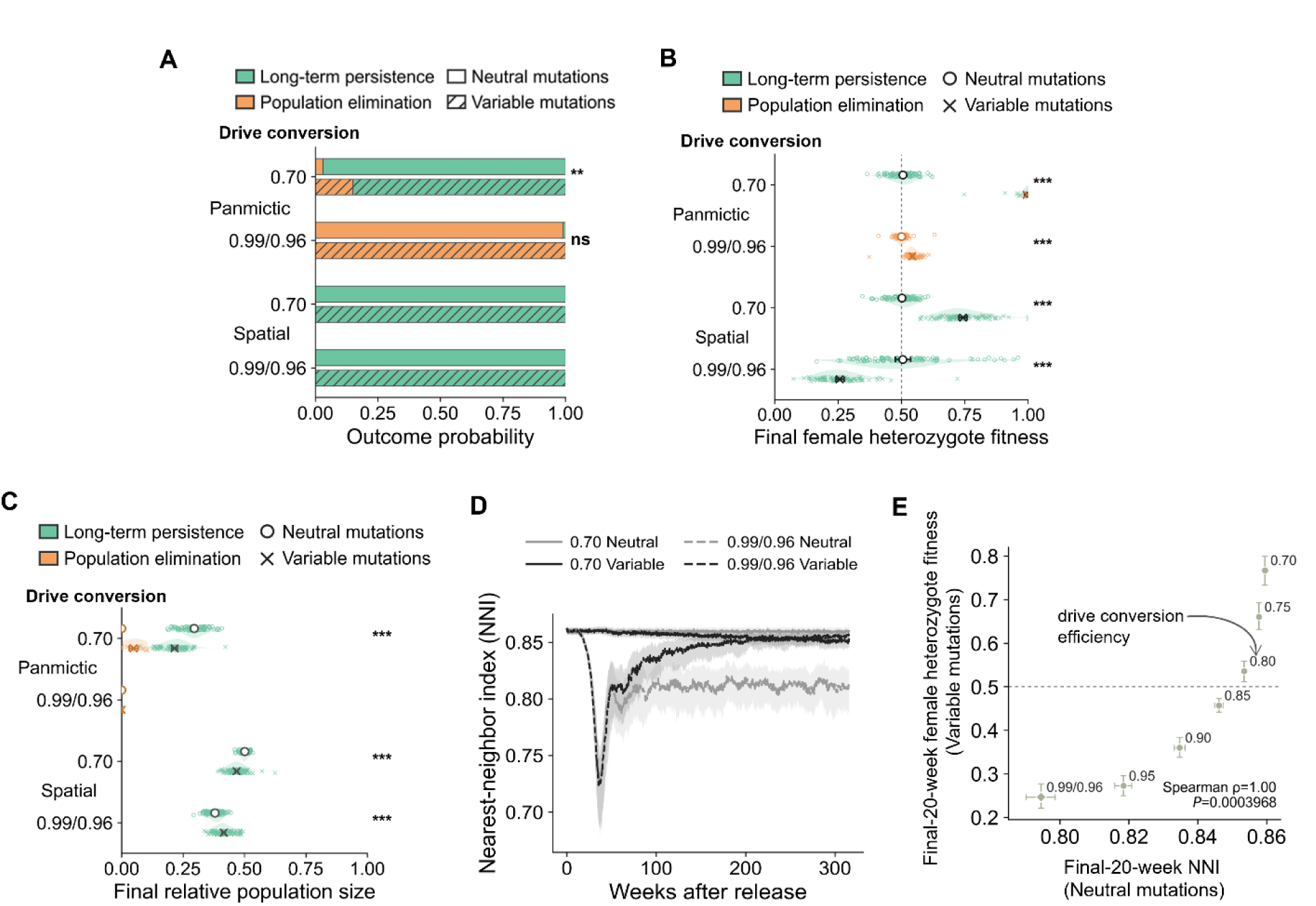
Mutations affecting female drive fitness evolve in response to population suppression gene drive. (**A**) Outcome composition across all simulations. Wilson 95% confidence intervals for outcome probabilities are reported in the Source Data. Th comparison was evaluated using a two-sided Fisher exact test (*P* = 0.014018). Final female heterozygote fitness in independent variable-mutation and neutral-mutation simulations. 0.99/0.96 refers to female and male drive conversion efficiency, respectively. (**B**) The endpoint for each simulation was defined as the adult-population-size-weighted mean of the final 20 weeks for population-level average fitness. Half-violins and small points show replicate-level distributions. Large dots indicate means, and horizontal error bars show 95% confidence intervals. The dashed line indicates the initial fitness of 0.5. Variable and neutral mutation groups were compared separately within each drive regime using two-sided independent two-sample permutation tests. Exact sample size, mean differences, and raw and adjusted *P* values are reported in Table S2. (**C**) Average relative long-term population size of the last 20 weeks compared to initial level. Plotting conventions and bootstrap intervals are as above. (**D**) Spatial NNI trajectories among simulations ending in long-term persistence. Solid lines show medians and shaded ribbons indicate the 25th–75th percentile interval. (**E**) Parameter-level association between the neutral-control mean NNI over the final 20 weeks and the corresponding mean final female heterozygote fitness in functional simulations across seven conversion settings. Each group contained 100 simulations. The association between final-20-week NNI of neutral control and final female heterozygote fitness in simulations with variable mutations was evaluated using a two-sided exact Spearman permutation test ( *ρ*=1.0, *P* = 0.000397). ns *P* ≥ 0.05; \**P* < 0.05; \*\**P* < 0.01; \*\*\**P* < 0.001.

Across all simulations, the high efficiency drive ended in population elimination in the panmictic model. The low efficiency drive failed to eliminate the population in controls, but could eliminate the population a fraction of simulations (7/100) when mutations could affect fitness (two-sided Fisher exact contrast, unadjusted *P* = 0.014). The spatial regimes were dominated by long-term persistence (Figure 3**A**). Variable fitness mutations evolved towards higher fitness in the low efficiency drives and in the high efficiency drive in the panmictic model (though elimination occurred quickly regardless). However, fitness evolved to lower levels for the high efficiency drive in the spatial model, whereas neutral-effect controls remained close to their baseline values (two-sided permutation tests, all adjusted (*P* < 0.001); Figure 3**B**). Long-term population sized was increased by 30% with the evolution of fitness mutations in the spatial high-conversion regime, but it was reduced by evolution in the low efficiency drives for both models (Figure 3**C**). These results show that fitness-mutation evolution was regime dependent and reversed direction in the high-conversion spatial regime, similar to embryo resistance evolution.

Spatial NNI trajectories likewise differed between variable mutations and control simulations for the high efficiency drive in the spatial model. In all simulations, the NNI increased with time, eventually reaching the value of the low efficiency simulations, indicating that chasing had likely completely ceased, with the population ending in an even equilibrium state (Figure 3**D**).

Adding several additional values of drive conversion efficiency (0.70, 0.75, 0.80, 0.85, 0.90, and 0.95, equal between females and males) where germline resistance was set to 0.5(1 - drive conversion) (other parameters were as described above), we examined the association between evolutionary responses and baseline spatial dynamics. All simulations in spatial models still ended in long-term persistence, and neutral-control mean NNI was positively correlated with fitness (Spearson’s *ρ* = 1.0, *P* = 0.000397; Figure 3**E**). Parameter settings producing more equilibrium-like outcomes (as opposed to chasing) in the neutral mutation model were associated with higher final attributes in the variable model. Together, female fitness evolution was regime dependent in the same manner as embryo resistance.

### Realized standing variation predicts evolutionary responses across genetic architectures

We next asked whether the amount of standing trait variation actually present at the time of drive release predicted subsequent evolutionary responses and whether genetic architecture provided additional explanatory information beyond realized average trait value from standing variation (**Figure 4**). For each mutable attribute, we examined two parameterization paths: one holding the number of mutation loci (*m*=50) constant while varying the effect-size scale (*b*), and the other holding *b* at its trait-specific minimum while varying *m*. We fitted a full interaction model relating each trait endpoint to trait standard deviation, parameterization path, and their interaction. Two joint Wald tests assessed the initial trait standard deviation effect and the additional genetic-architecture effect, with *P*-values Holm-adjusted separately across the three traits (see methods for details).

**Figure 4.**
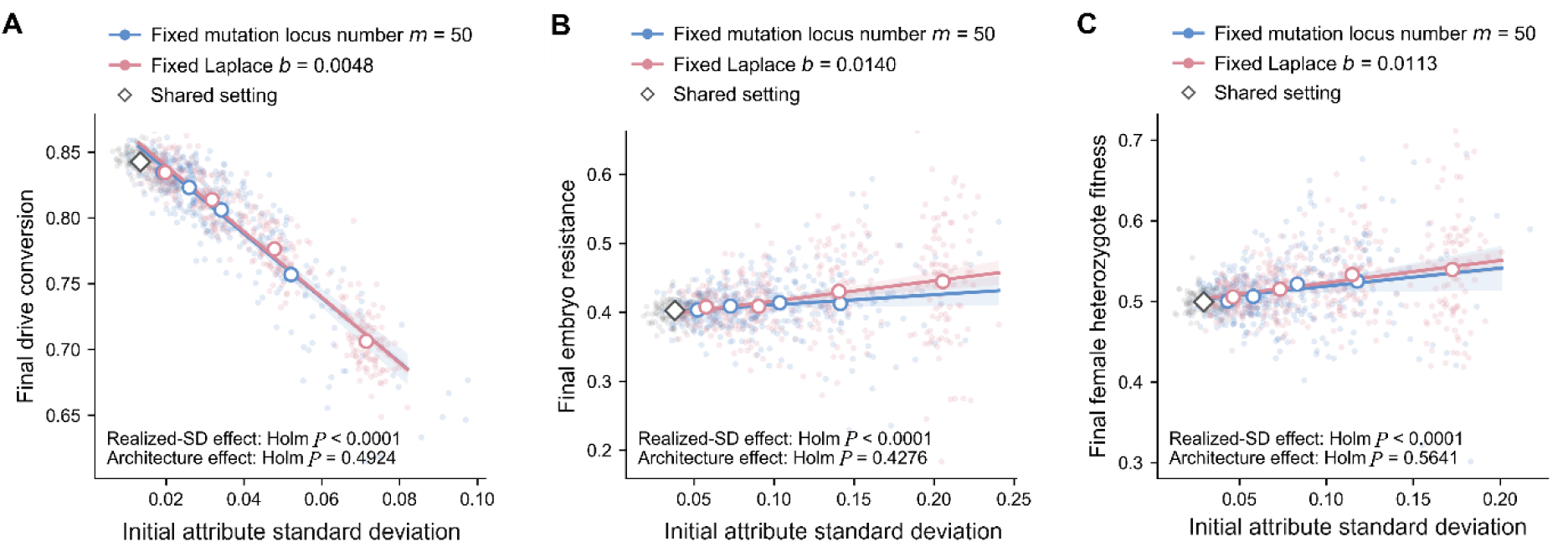
Final mutation attributes based on different initial distributions of standing trait variation. Fixed background parameters were the spatial model, female heterozygote fitness 0.5, and other parameters as default in Table S1. Charts show (**A)** final drive conversion, (**B)** final embryo resistance, and (**C)** final female heterozygote fitness. Each small point represents one independent simulation, and each parameter setting contained 100 simulations. Blue denotes the fixed-m path (*m* = 50), orange denotes the trait-specific fixed-b path, and open circles denote parameter-setting means. The shared *m* = 50, *b* = *b*min setting is shown once in gray and was excluded from both path-comparison arms. Realized pre-release attribute standard deviation is the mean adult attribute standard deviation over weeks 11-20 before release. Solid lines and shaded bands show estimates and 95% confidence intervals from the full interaction model.

Realized pre-release standing variation was significantly associated with the final evolutionary endpoints of all three mutable attributes (**Figure 4**). Greater realized initial attribute standard deviation was associated with lower final drive conversion, with nearly identical negative slopes along the fixed-*m* and fixed-*b* paths (−0.0245 and −0.0250 per 0.01-unit increase in attribute standard deviation, respectively; Holm-adjusted *P* < 10⁻³⁰⁰; **Figure 4A**). In contrast, embryo resistance and female heterozygote fitness increased with realized standard deviation along both paths, although their associations were weaker than that of conversion (Holm-adjusted *P* = 2.18 × 10⁻⁷ and 2.13 × 10⁻⁷, respectively; **Figure 4B-C**). Despite these consistent associations, neither the joint genetic-architecture effects (Holm-adjusted *P* = 0.4924, 0.4276, and 0.5641 for conversion, embryo resistance, and female heterozygote fitness, respectively) nor the individual slope differences between paths were statistically significant. These results indicate that realized standing variation at drive release was a consistent predictor of subsequent evolutionary responses, whereas the two genetic-architecture parameterization paths provided no detectable additional explanatory information after accounting for realized standard deviation. Together, these results show that pre-release standing variation of traits was positively associated with the magnitude of subsequent evolutionary change across all three drive properties. In contrast, we found no statistically significant evidence that the two genetic-architecture parameterization paths (more mutations or larger effect size per mutation) provided additional explanatory information. These findings identify realized standing variation as a consistent predictor of evolutionary response under the architectures examined, while not establishing that genetic architecture has no effect (which may become more important with very small numbers of mutations) or that the two paths are exactly equivalent.

### Suppression drive changes dispersal evolution

Previous studies of chasing indicated that dispersal can be a critical parameter for the outcome (Champer et al., 2021; Champer et al., 2022; Combs et al., 2026; Liu and Champer, 2022; Liu et al., 2023; Paril and Phillips, 2022; Zhu and Champer, 2023), and it is reasonable to expect that dispersal would vary in natural populations. Thus, we examined the effect of genetic variation for dispersal in suppression drive settings. It was necessary to impose a fitness cost for high dispersal to balance the initial state in a no-drive scenario (see methods). We found that ultimate outcome composition did not differ between evolving and fixed-dispersal populations at drive conversion efficiencies of 0.75 and 0.85, whereas significant differences emerged at 0.95 and 0.99 ( Figure 5**A**). At 0.95, population elimination occurred in 7% of evolving populations compared with none of the fixed-dispersal controls. At 0.99, elimination occurred in 18% versus 1%, respectively, while drive loss occurred in 6% versus 1% (drive loss is often correlated with successful elimination outcomes (Liu et al., 2023)). Thus, allowing dispersal evolution altered the distribution of system outcomes at the higher conversion efficiencies.

**Figure 5.**
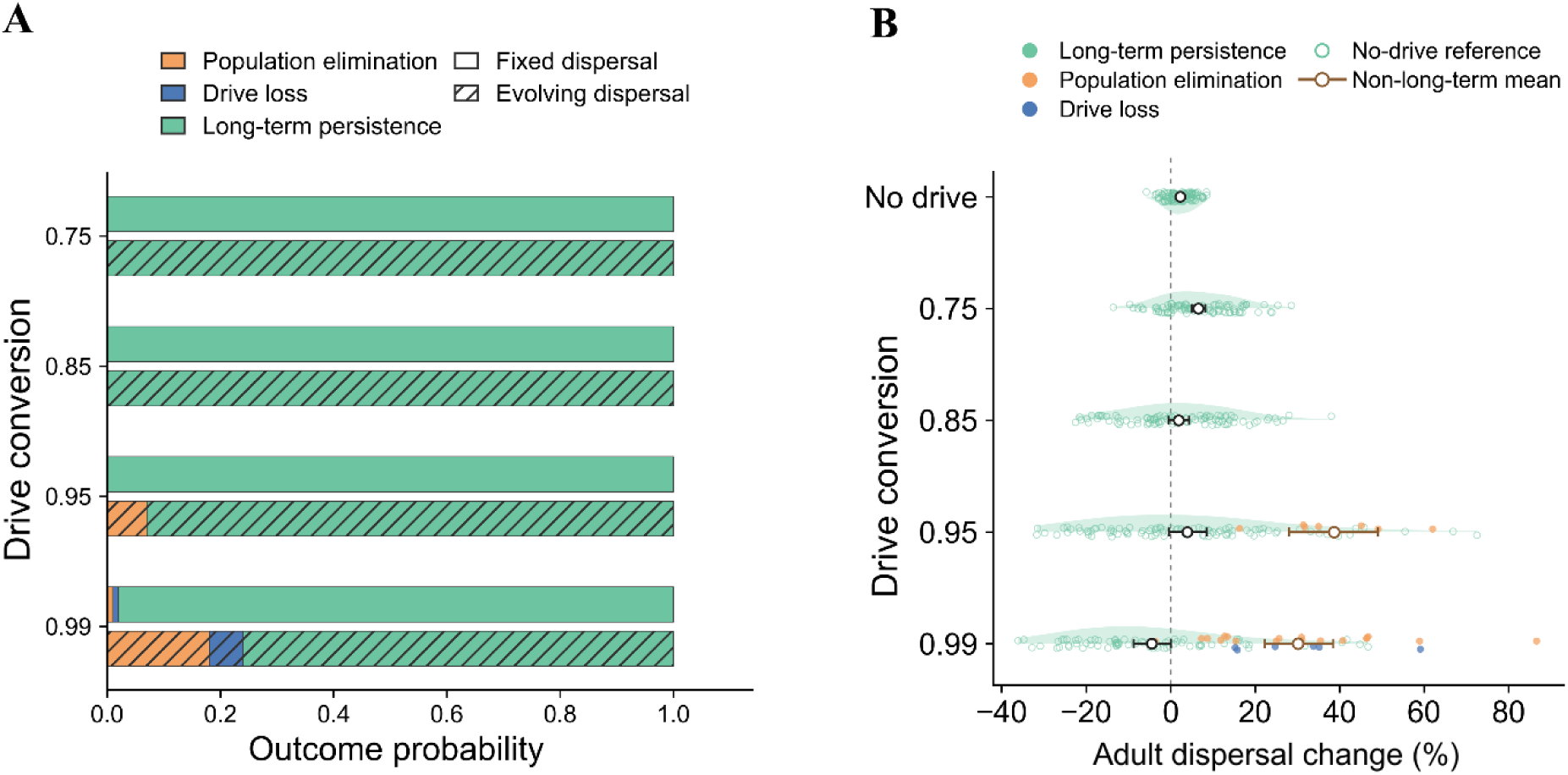
Effects of standing genetic variation that influence dispersal on suppression drive outcomes. Evolving-dispersal simulations were compared with fixed-dispersal controls. The pre-release baseline was week 100, and drive release occurred at week 101. Each parameter set had 100 replicates. (**A**) Outcome composition across all simulations. Overall outcome composition was compared between dispersal treatments within each conversion setting using a two-sided Fisher–Freeman–Halton exact test, with Holm correction across four comparisons. (**B**) Cumulative adult dispersal change among evolving populations, with no-drive populations providing a background reference. Change was calculated based on the adult-population-size-weighted average of the final 20 weeks relative to mean dispersal at week 100. Half-violins and small points show replicate distributions. Large dots indicate arithmetic means, and horizontal error bars show 95% confidence intervals. The vertical dotted line indicates zero change. Each drive group was compared with its origin-seed-matched no-drive counterparts using a two-sided paired t-test. These four comparisons were jointly Holm-adjusted. Significance symbols indicate Holm-adjusted *P* values: ns *P* ≥ 0.05; \**P* < 0.05; \*\**P* < 0.01; \*\*\**P* < 0.001. Exact sample sizes, effect estimates, and raw and adjusted *P* values are provided in the Source Data.

We next quantified cumulative dispersal changes. Dispersal change was calculated as the percentage change in mean dispersal over the final 20 weeks relative to the pre-release baseline. No-drive populations showed a small positive change, providing a reference for background evolution. At a conversion efficiency of 0.75, evolving drive populations exhibited a greater increase than their matched no-drive counterparts (Figure 5**B**), perhaps because selection pressure favored higher dispersal when density was reduced, changing the equilibrium value that was better balanced for a no-drive scenario. The corresponding contrasts were not significant at 0.85 or 0.95 after multiple-comparison correction, though the standard deviation of dispersal change further increased from the no-drive scenario. At 0.99, however, coexisting populations showed a significant negative average dispersal change. In both 0.95 and 0.99, simulations that resulted in population elimination had a substantially increased dispersal. Thus, although the highest conversion efficiencies may increase the probability of elimination or drive loss by stochastically reducing dispersal from population bottlenecks, populations that ultimately maintained long-term coexistence often exhibited a decline in mean dispersal ability.

### Recessive deleterious variation deepens the population bottleneck and destabilizes persistence

Inbreeding and deleterious recessive mutations are common in most species. Previous gene drive studies have shown that inbreeding difficulties could assist the drive, but these used simplified models (Bull, 2016; Bull et al., 2019; Champer et al., 2021). We thus assessed such genetic variation with explicit mutations. Before drive release, panmictic and spatial populations were simulated for 6,000 weeks to approach mutation–selection–drift equilibrium. The virtual inbreeding load and number of deleterious mutations per individual subsequently remained stable (**Figure S3**). Panmictic individuals carried more deleterious mutations than spatial individuals, despite the higher mean fitness of panmictic populations. This was likely because random inbreeding was higher in spatial populations. Thus, the two population models differed in both the inbreeding load and potential consequences of recessive deleterious alleles before release.

We then released suppression drives into populations simulated with or without recessive deleterious variation. Populations carrying recessive deleterious variation reached lower minimum post-release population size relative to their own pre-release baselines in most scenarios, but no difference was detected in the high drive efficiency panmictic scenario where population elimination was achieved (Figure 6**A****, Figure S4**). The observed elimination fraction was higher in load-bearing populations in the low drive efficiency panmictic scenario (Figure 6**B**). The spatial simulations retained long-term persistence, irrespective of recessive deleterious variation.

**Figure 6.**
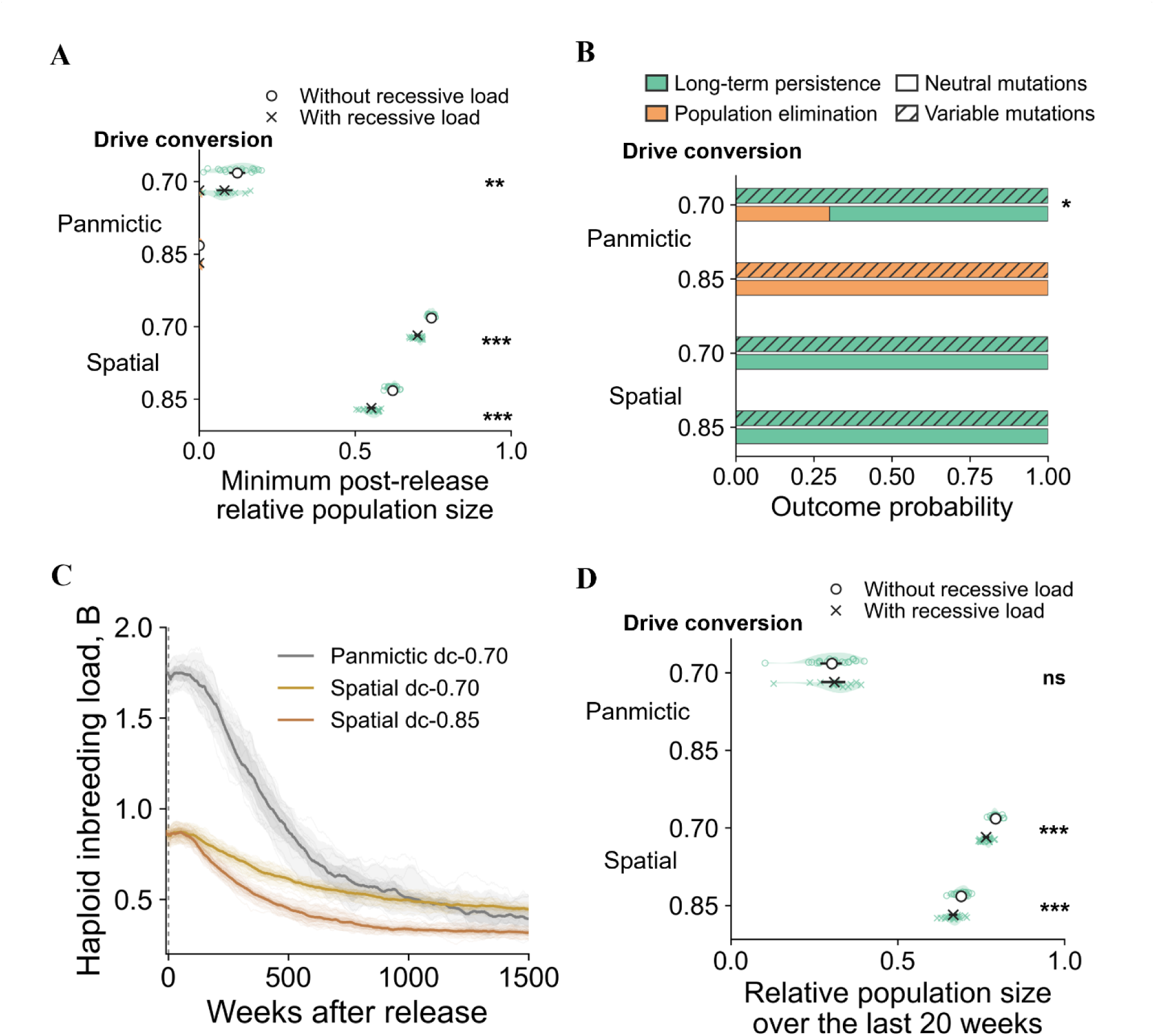
Recessive deleterious variation deepens the post-release bottleneck and alters simulation outcomes. (**A**) Minimum post-release relative population size for simulations with and without recessive deleterious variation. Half-violins and small points show replicate-level distributions. Large dots indicate means, and horizontal error bars show 95% confidence interval. Each condition and treatment contained 20 simulations. The corresponding valley week is reported in the Source Data. (**B**) Outcome composition across all simulations. Wilson 95% confidence intervals for outcome probabilities are reported in the Source Data. Two-sided Fisher exact tests for elimination in Panmictic-0.70 scenario is reported (*P* = 0.020196). (**C**) Release-aligned virtual haploid inbreeding-load trajectories for load-present simulations that reached long-term persistence. Thin lines are individual simulations, thick lines are across-simulation medians, dark ribbons are the central 50% interval, and light ribbons are the central 90% interval. Panmictic–0.85 is omitted because no load-present simulation reached long-term persistence. (**D**) Relative long-term population size of the last 20 weeks. Plotting conventions and bootstrap intervals are as above.

Among load-bearing replicates in which both the population and drive persisted, virtual haploid inbreeding load declined after release in all three scenarios with long-term persistence (Figure 6**C**). Most of the decrease occurred during the first several hundred weeks, followed by a slower decline. This decline occurred despite continued natural formation of deleterious mutations and corresponded to a net reduction in the additional viability cost exposed by the virtual full-sib assay. The high efficiency panmictic scenario was not represented because it resulted in rapid population elimination. Consistent with this, the recessive mutation number per individual sharply declined, and the average viability of the population increased almost to pre-release levels at the end (**Figure S5**). Long-term relative population size remained lower in load-bearing spatial populations, whereas no difference was detected in the low efficiency panmictic scenario (Figure 6**D**). This shows a rapid loss of influence for standing recessive deleterious mutations. In spatial populations, relative long-term population size was generally lower with recessive deleterious variation than without it (Figure 6**D**). Thus, declining virtual inbreeding load was not accompanied by convergence to the no-load controls in all demographic settings.

Together, these results show that recessive deleterious variation amplified demographic vulnerability following drive release. It deepened the post-release population minimum and possibly reduced persistence. Much of the inbreeding load was subsequently purged from populations that recovered, concentrating its strongest effects in the bottleneck and outcome-determination phases.

## Discussion

Gene drive research has progressed from proof-of-principle homing constructs to laboratory cage suppression experiments (Du et al., 2024; Kyrou et al., 2018; Morianou et al., 2026; Xu et al., 2024). These studies demonstrate that high transmission can support population suppression. They also show that fertility costs, resistance formation and ecological context determine whether laboratory performance scales to population collapse. Most experimental and modelling studies consequently treat conversion efficiency, resistance, and drive fitness as measured properties of a construct. Our results address a complementary question: whether those properties remain evolutionarily stable when a fixed construct encounters standing variation in the surrounding genome that can affect them.

Most studies of gene drive evolution have focused on resistance mutations at the target site. Functional resistance alleles arising from standing variation or cleavage-associated repair can spread and reduce drive performance (Champer et al., 2018; Champer et al., 2020; Hammond et al., 2017; Noble et al., 2017; Unckless et al., 2017). Weakly deleterious natural resistance variants may remain a substantial source of escape. Empirical work in genetically diverse *Drosophila* backgrounds further shows that conversion efficiency and embryo resistance formation are heritable quantitative traits without one common large-effect locus (Champer et al., 2017; Champer et al., 2019). Thus, it remains plausible that in some scenarios, the evolution of genomic background in response to suppression may be important.

When might this be an important consideration? In terms of magnitude, we used conservative estimates for the effects of genomic variation (see methods), and thus our study should be interpreted as a general framework for what direction evolution is likely to take rather than a quantitative prediction of specific scenarios, which depend highly on species-specific background. From another perspective, we modeled genomic effects as additive, which is likely a suitable approximation when variation can be important. However, the effects of genomic variation on drive parameters may be nonlinear, reflecting underlying molecular processes such as Cas9 cleavage and DNA repair. The importance of cleavage timing and Cas9 activity saturation has previously been demonstrated in models of homing drives with multiplexed gRNAs (Champer et al., 2020). Based on such a paradigm, the effects of genomic variation might be considerably less important for parameters near extreme ends of the range. Thus, *Anopheles gambiae* suppression drives with ∼98% drive efficiency and ∼10% embryo resistance (Champer et al., 2022; Kyrou et al., 2018) would not suffer major effects from evolution of standing variation on these parameters. This was borne out in initial tests of drive conversion efficiency of a similarly effective drive in this species (Pescod et al., 2023). However, the female heterozygote fitness of the *dsx* suppression drive was at an intermediate level (Champer et al., 2022; Kyrou et al., 2018) and is thus potentially more likely to vary between individuals with different genomic backgrounds. Similarly, a toxin-antidote drive in *Drosophila melanogaster* with high embryo resistance did not have this parameter change in genomic background (Oberhofer et al., 2019), while a homing drive had considerable variation in this parameter when it was at an intermediate level in a lab strain (Champer et al., 2017; Champer et al., 2019). Given the challenge of achieving high drive conversion in non-*Anopheles* species (Chen et al., 2026; Zhang et al., 2026), it is possible that genomic variation may play an important role in the consideration of several future suppression drives.

Our simulations generalize genomic resistance beyond the target sequence, and strictly speaking, even beyond the concept of “resistance”. The genetic background can modify transmission, resistance formation and drive fitness, and these components need not share one evolutionary response. Across the tested regimes, mutation classes differed in evolutionary direction. Conversion mutations consistently reduced transmission when sufficient post-release selection accumulated, reducing drive performance. Embryo resistance and drive fitness mutations instead improved drive performance, except perhaps in chasing scenarios, where they too could reduce drive effectiveness. These contrasts indicate component-specific and regime-dependent evolutionary responses. They do not support a single genomic response to drive release.

Viewed component by component, the bidirectional responses may arise because each mutation changes a different balance of immediate reproductive effects and delayed lineage consequences. Reduced conversion can protect background lineages from drive-associated genetic load, making attenuation advantageous across a broad range of contexts. Embryo resistance creates a more contingent trade-off between the short-term cost of producing nonfunctional resistant offspring and the longer-term benefit of avoiding repeated conversion. Fitness mutations likewise balance a direct benefit in fertile drive heterozygotes against costs incurred when mutation-bearing descendants remain associated with drive backgrounds. The observed sign changes are consistent with these competing effects shifting across regimes, although the present design does not isolate any single term.

One possible explanation is that embryo resistance changes descendant-lineage persistence rather than providing a uniform immediate advantage. Nonfunctional resistance can contribute to sterile females when both target-site copies are disrupted, creating a short-term reproductive cost. Resistant alleles transmitted through males, or through genotypes that retain reproductive function, are nevertheless refractory to subsequent conversion. Under repeated drive exposure, this escape from conversion could increase the relative persistence of resistance-bearing lineages compared with lineages that repeatedly re-enter drive-associated, fertility-reducing genotypes. The balance may therefore favor lower embryo resistance in less exposed regimes but shift towards higher resistance during prolonged chasing, when lineages tend to have more inbreeding, preserving the longer-term advantage of having some nonfunctional resistance alleles present.

Female heterozygote fitness evolution can be interpreted similarly through a parallel balance of genotype-specific benefits and lineage-level costs. A mutation that increases somatic fitness can provide an immediate reproductive advantage when it is expressed in fertile female drive heterozygotes. The same mutation may, however, become associated with drive-bearing descendants, including sterile female drive homozygotes that remove mutation copies from future reproduction. The net response therefore depends not only on drive-heterozygote frequency, but also on associations between mutation alleles/drive genotypes and on the population structure of their descendants. This balance could favor higher drive fitness in well mixed regimes such as a panmictic population or spatial populations with a homogenous equilibrium, but lower fitness in a chasing environment.

These patterns do not demonstrate convergence towards a universal optimum. Establishing such an attractor would require a continuous drive-strength gradient and direct evidence that evolutionary changes converge from both sides. At the system level, these contrasting responses could buffer drive pressure asymmetrically, with perhaps more routes available to attenuate a strong drive than to strengthen a weak one in a chasing scenario. This is a hypothesis about patterns across the tested regimes, not a prediction that all drives converge to a shared intermediate optimum.

These results are consistent with a reciprocal eco-evolutionary feedback between mutation evolution and drive-induced population dynamics. Standing mutations altered drive performance and the subsequent demographic or spatial state of the population. Those realized states also defined the context in which further selection occurred. Covariation with drive frequency and NNI is therefore consistent with a coupled feedback process, not a one-way pathway controlled by one demographic variable. Spatial regimes did not merely change endpoint distributions (Bull et al., 2019; Champer et al., 2021; North et al., 2020). They were also the contexts in which embryo resistance and fitness responses changed direction. This does not identify chasing as the sole causal mediator, but it shows that spatial dynamics and background evolution cannot be forecast independently. The simulations establish regime-dependent responses and functional feedback, while the identity of the mediating state variables remains unresolved.

Our analyses further show that the evolutionary potential of standing variation did not show significant differences among different genetic architectures. Genetic architecture determines how phenotypic variation is distributed among loci and can therefore influence evolutionary trajectories even when populations have similar initial genetic variance (Hansen, 2006; Hayward and Sella, 2022). Consistent with this view, previous work has shown that effect-size distributions, demographic context, redundancy, allele frequencies, and linkage disequilibrium can all affect the rate and predictability of polygenic adaptation (Schlötterer, 2023; Stetter et al., 2018). Our simulations extend this principle to the evolution of drive-modifying traits. At comparable levels of realized standing variation (more mutations of smaller effect or vice versa), we found no detectable differences in outcomes between the two parameterization approaches. This result does not establish equivalence between the paths or exclude effects of genetic architecture outside the parameterizations examined.

Allowing dispersal evolution from standing variation altered suppression drive outcomes at higher conversion efficiencies, suggesting feedback between dispersal traits and demographic dynamics. Initially, we expected that during chasing, high-dispersal wild-type individuals could potentially gain an advantage by avoiding the drive, thus increasing population average dispersal and thus potentially leading to better future drive outcomes. This did not seem to occur consistently, perhaps due to faster drive elimination immediately after this evolved in part of the population. However, dispersal still played an important role in outcomes. At a drive conversion efficiency of 0.99, populations maintaining long-term coexistence showed reduced mean dispersal, whereas simulations ending in elimination or drive loss generally showed increased dispersal. These contrasting patterns are compatible with population bottlenecks generating different evolutionary trajectories, with drift and possibly selection acting on the remaining variation. However, the coexistence estimates are conditional on the final outcome and cannot establish that prolonged chasing consistently favors lower dispersal. Moreover, the different endpoint windows and continued evolution after possible drive loss limit causal comparisons across outcomes. Our results reveal an association between dispersal change and drive fate, while leaving the relative contributions of genetic drift, directional selection and outcome-dependent sampling unresolved. Recessive deleterious variation followed a distinct feedback pathway. Rather than changing a drive-related trait directly, it deepened post-release demographic decline and weakened recovery from low density. These results are consistent with inbreeding research showing a decrease in long-term per-capita population growth and an increase in contraction-related risks (Charlesworth and Willis, 2009; Kyriazis et al., 2021). The subsequent reduction in deleterious load among recovered populations indicates that this feedback was strongest during the transient decline. This mechanism extends gene drive modelling by treating recessive load as a dynamic genetic state rather than a fixed inbreeding-depression penalty (Beaghton and Burt, 2022; Bull et al., 2019; Champer et al., 2021). A population can carry masked deleterious variation before release without an obvious effect. A drive-induced decline can expose substantial realized load when homozygosity increases. Accurate forecasts may also require information regarding mutation dominance, homozygous load, and low-density growth rate.

Together, the mutation and recessive load analyses identify two routes through which genomic background enters drive forecasts. First, standing variation makes drive-performance components evolutionarily labile. Second, hidden recessive load makes demographic recovery state-dependent during population decline. Figure 7 integrates these routes within one eco-evolutionary framework. The genomic background is therefore part of the dynamical system, rather than a neutral scaffold surrounding a fixed drive cassette.

**Figure 7.**
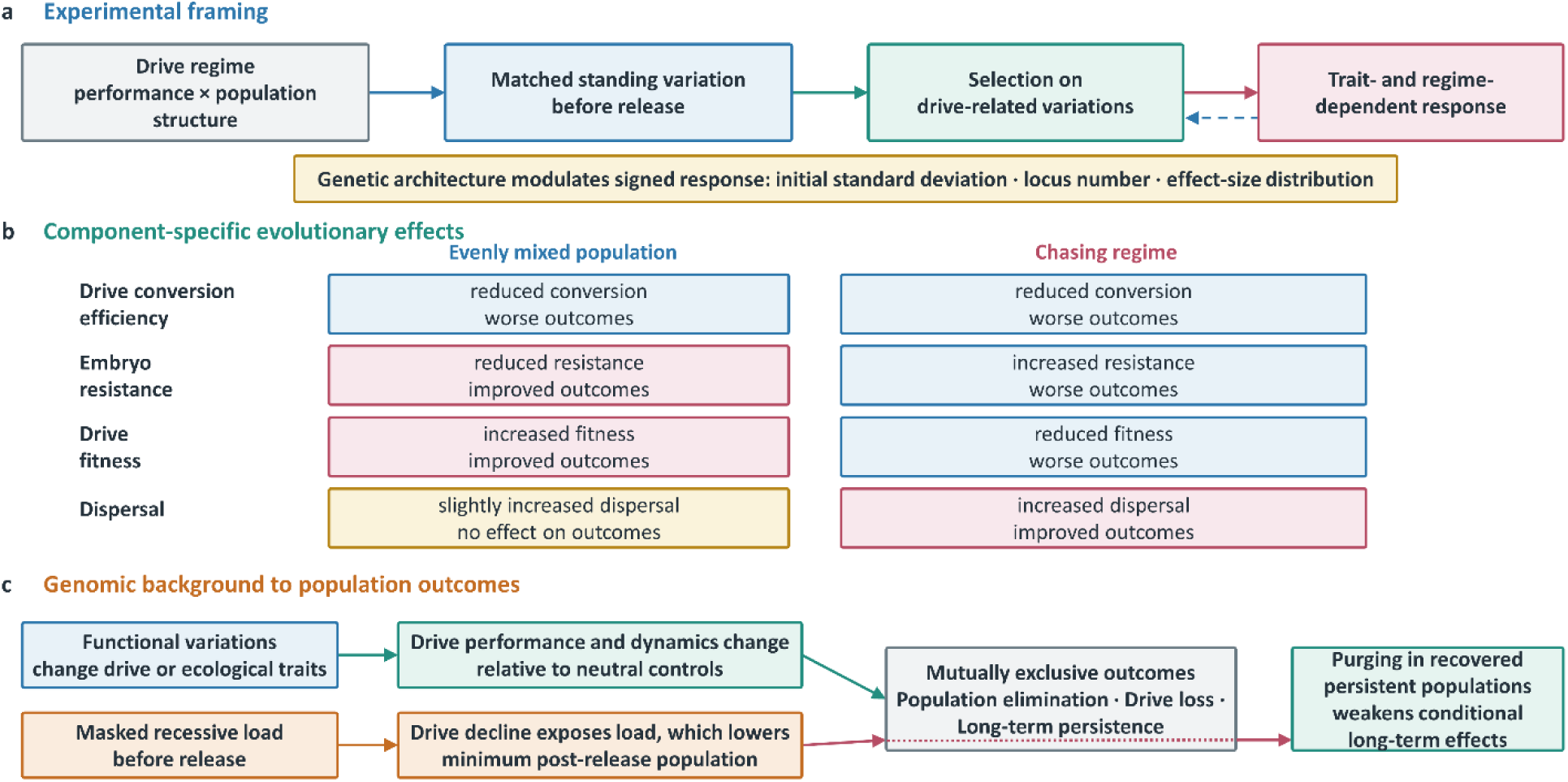
Conceptual synthesis of standing genetic variation interactions with suppression gene drive. (**A**) Drive performance and population structure define experimental drive regimes in which standing variation is selected for. (**B**) Drive conversion efficiency mutations show a directionally robust anti-drive response, whereas embryo resistance and fitness mutations reverse direction across regimes. Evolution of dispersal may also favor successful elimination outcomes in a chasing regime. (**C**) Variable mutations alter drive traits and subsequent dynamics relative to neutral controls, whereas recessive deleterious variation follows a separate demographic pathway through load exposure, a lower post-release population minimum, outcome redistribution, and purging among recovered persistent populations.

There are several additional conditions we did not model that may further influence a gene drive system acting in an environment where genomic variance affects performance. Several genomic features were deliberately simplified. One limitation is that sex-linked mutational variation was not simulated. The non-recombining region of the *Anopheles* Y chromosome is highly repeat-rich but gene poor and may thus have only modest effects (Hall et al., 2016). The X chromosome also contains fewer annotated genes than the autosomes, but still has large numbers of genes with a unique inheritance pattern (Holt et al., 2002; Kojin et al., 2022) that could respond differently to sexually asymmetric drive pressure, as seen in female fertility homing suppression drives. Hemizygosity in males is expected to expose recessive X-linked alleles to selection (Charlesworth and Charlesworth, 2010; Ellegren, 2011), and the X chromosome can also harbor variants with sex-specific fitness effects (Innocenti and Morrow, 2010). Sex-linked mutations may therefore follow different dynamics from autosomal mutations.

Another limitation is that the genomic organization of mutation loci was represented by random distributions across chromosomes. Although our simulations explicitly incorporated chromosomal positions and recombination, natural genetic architectures are more complex because recombination rates are heterogeneous across genomes, and linked variants can constrain the independent response of alleles to selection (Hill and Robertson, 1966). The evolutionary potential of complex traits depends not only on the number and effects of contributing loci, but also on their genomic organization and linkage relationships (Barton et al., 2002). Future studies incorporating empirically informed recombination landscapes and non-random genomic architectures will help determine how genome organization influences the evolvability of gene drive mutations.

Several simplifying assumptions also affect how these results should be interpreted. Mutation alleles were modelled as neutral with respect to fitness components other than the trait they modified, and the initial allele-frequency spectrum is consistent with the neutral theory of molecular evolution (Ewens, 2004; Tajima, 1989). However, pleiotropic fitness effects would change pre-release frequencies and subsequent selection. Linkage to beneficial or deleterious alleles could also redirect the response (Hill and Robertson, 1966). Mutation loci were not allowed to interact epistatically in our model. Empirical work in *Drosophila* has reported departures from independent multi-locus effects (Simmons and Crow, 1977; Stoffel et al., 2021), although the magnitude and direction of epistasis are not consistent across datasets (Temin et al., 1969). In addition, we modelled fully recessive mutations (h = 0) with multiplicative effects on survival, without a distribution of dominance coefficients, epistasis (Hedrick and Garcia-Dorado, 2016; Rashid et al., 2022) or effects on fecundity, mating success or dispersal survival (Harrisson et al., 2019). However, deleterious alleles with appreciable dominance are common and more efficiently removed from populations (Mukai et al., 1972) and may thus be of lower importance for gene drive scenarios.

Several additional limitations arise from the regime-level design. Conversion and population structure jointly changed drive frequency, genotype composition, population size and spatial dynamics, so the current comparisons cannot isolate NNI, drive frequency, or population size as a single mediator. Continuous post-release summaries are conditional on long-term persistence, and only selected mutation classes, genetic architectures and standing variation present before release were examined. It is also possible that variation in baseline drive performance could substantially increase or decrease observed outcomes based on variable mutations, especially if initial performance was closer to a “borderline” outcome regime compared to our default parameters.

Overall, our results suggest that suppression drives, at least in some scenarios, should be treated as evolving eco-genetic systems rather than engineered elements with permanently fixed performance. Standing variation can expose evolutionary change across drive transmission, resistance formation, and fitness, while hidden recessive load can reshape demographic recovery. The resulting feedback is component-specific and asymmetric. Incorporating these processes should improve theoretical forecasts without implying that all systems converge to a common evolutionary endpoint.

## Acknowledgements

This study was supported by the Center for Life Sciences and the National Natural Science Foundation of China (32270672, W2432018).

## Supplemental Information

**Table S1.** Default parameter values for simulations.

| Parameter | Value |
|---|---|
| Drive conversion | 0.85 |
| Germline resistance | 0.5*(1-drive conversion) |
| Embryo resistance | 0.4 |
| Female heterozygote fitness | 0.9 |
| Number of adult females | 10,000 |
| *β*<sub>*max*</sub> | 25 |
| Area | 1 |
| Average dispersal distance | 0.03 |
| Low-density growth rate | 6 |
| Foraging distance | 0.01 |
| Total gene element number | 10,000 |
| Mutation loci number | 100 |

### Supplemental Figures

**Figure S1.**
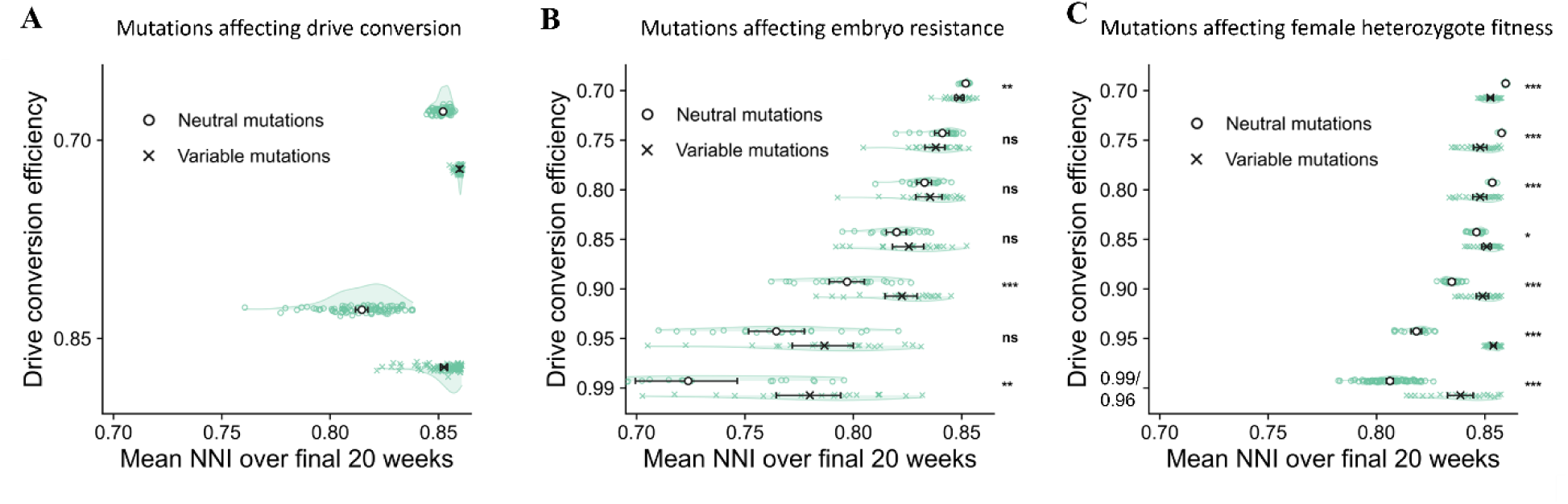
Spatial nearest-neighbor index with genomic variation affecting drive performance. Charts display nearest-neighbor index (NNI) in spatial simulations with genomic variation affecting (**A**) drive conversion efficiency, (**B**) embryo resistance and (**C**) female drive heterozygote fitness. 0.99/0.96 refers to female and male drive conversion efficiency, respectively. Each point is one simulation’s mean NNI over its final 20 weeks. Open circles denote neutral mutation controls and crosses denote variable mutation simulations. Half-violins show replicate distributions, and large black points and horizontal error bars show arithmetic means and bootstrap 95% confidence intervals. Each displayed group contains (**A**) 100 or (**B,C)** 20 simulations. Panel A complements the spatial trajectories in Figure 1D. Panels B and C complements the final NNI comparison between neutral and variable mutations of settings in Figure 2E and 3E, respectively.

**Figure S2.**
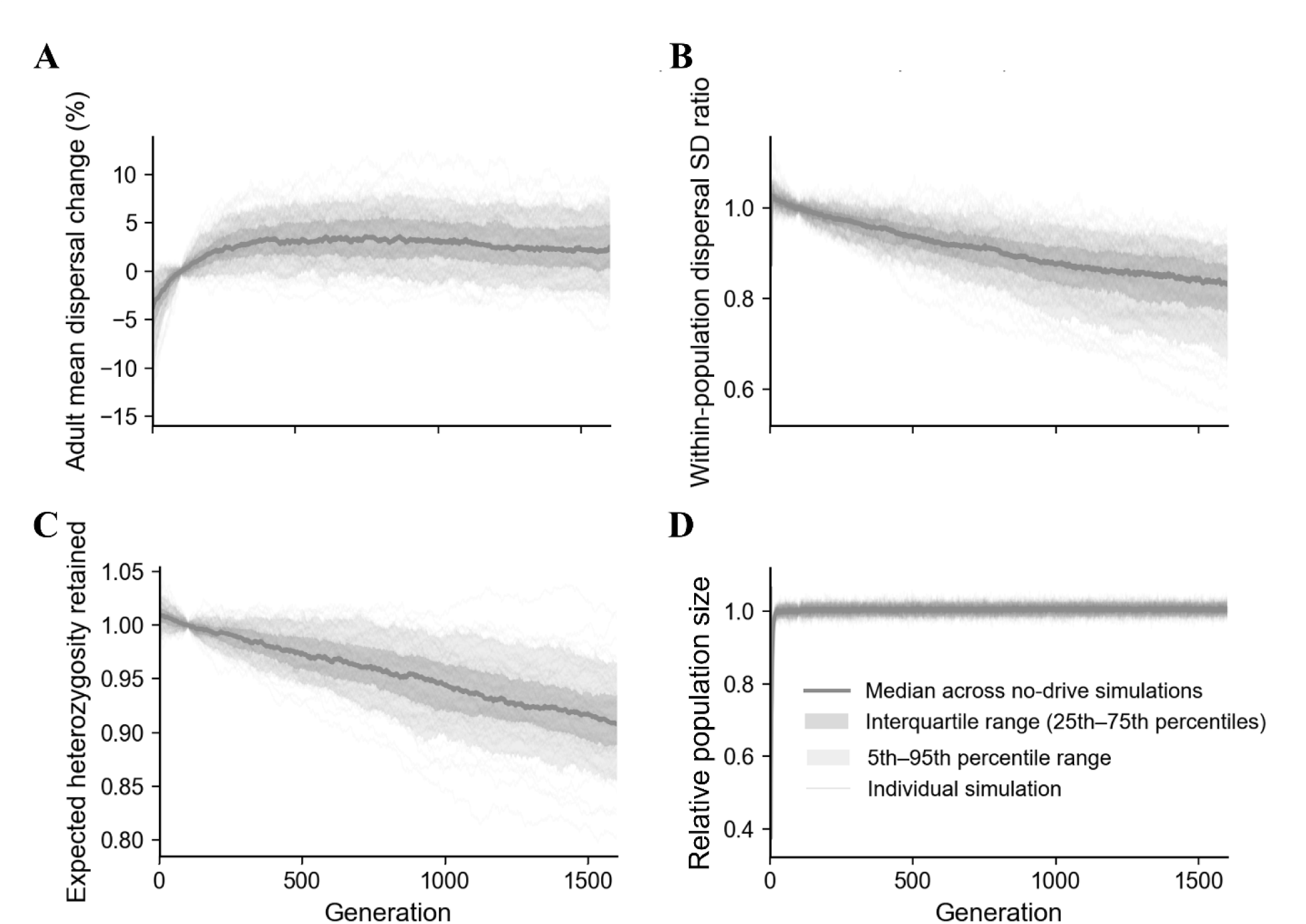
Dispersal variation and genetic diversity in the no-drive reference populations. (**A**) Percentage change in mean adult dispersal relative to the week-100 baseline. (**B**) Within-population dispersal standard deviation (SD) relative to its baseline. ( **C**) Expected heterozygosity retained. (**D**) Relative population size to carrying capacity. All panels include the initial 100-week transition and the subsequent observation period. Thin curves show individual simulations, thick curves show across-simulation medians, dark bands show the interquartile range (25th–75th percentiles), and light bands show the 5th–95th percentile range.

**Figure S3.**
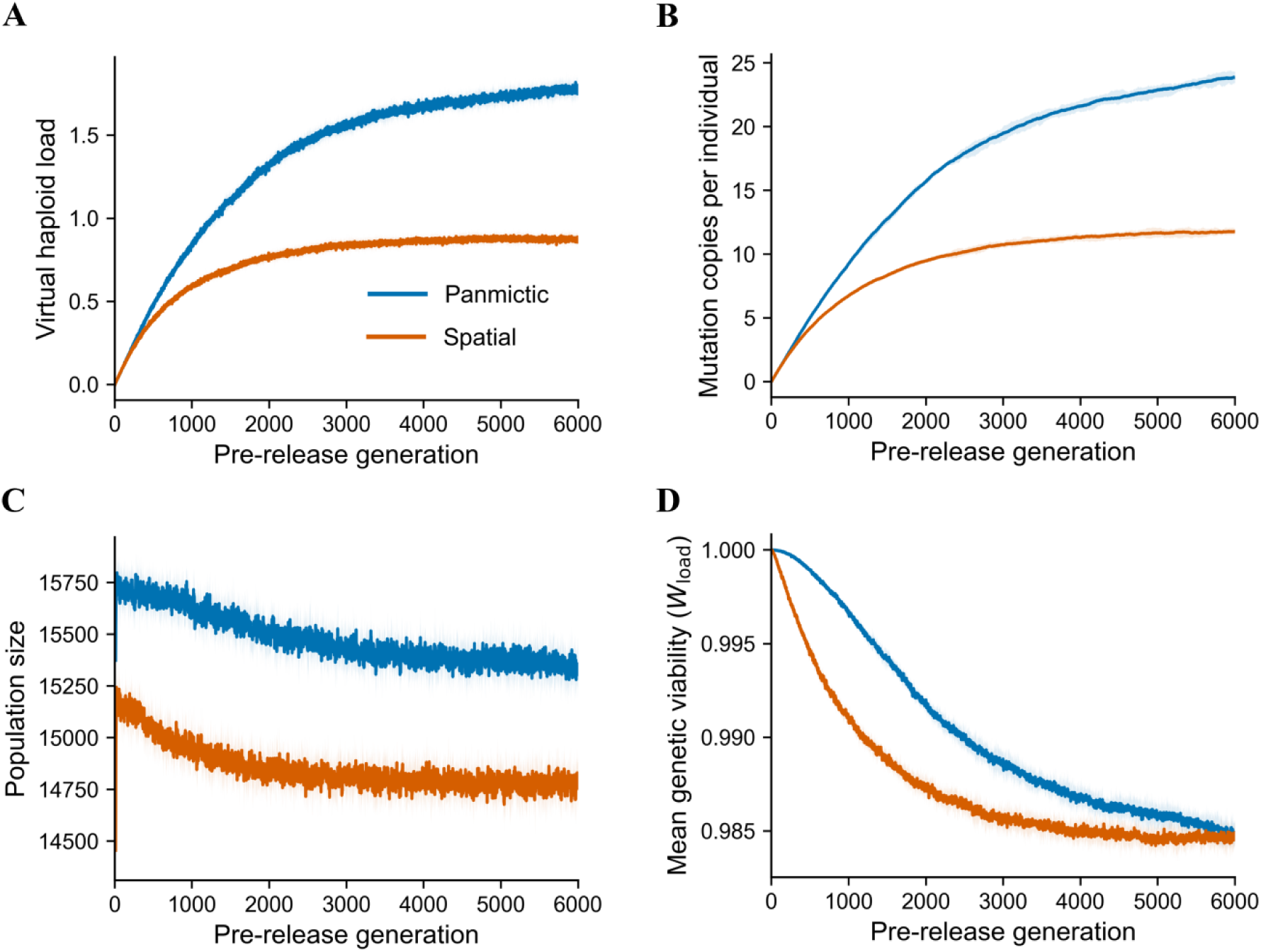
Establishment of recessive deleterious variation before drive release. Pre-release trajectories of (**A**) virtual haploid inbreeding load, (**B**) average deleterious mutation copies per individual, (**C**) population size, and (**D**) mean genetic viability. Lines show across-simulation median trajectories.

**Figure S4.**
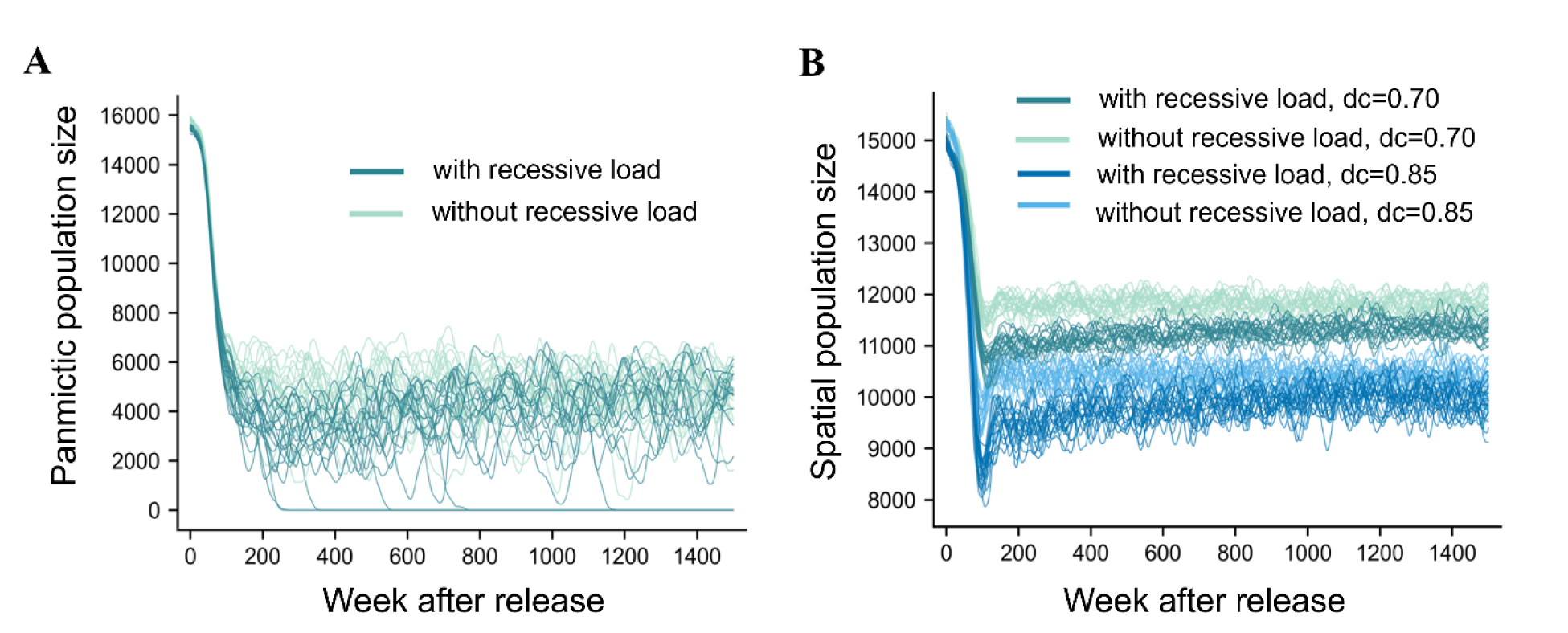
Population size trajectories with recessive deleterious variation. Graphs show number of individuals in (**A**) panmictic and (**B**) spatial populations. Each condition and treatment has 20 simulations. Each line represents a simulation. dc = drive conversion efficiency.

**Figure S5.**
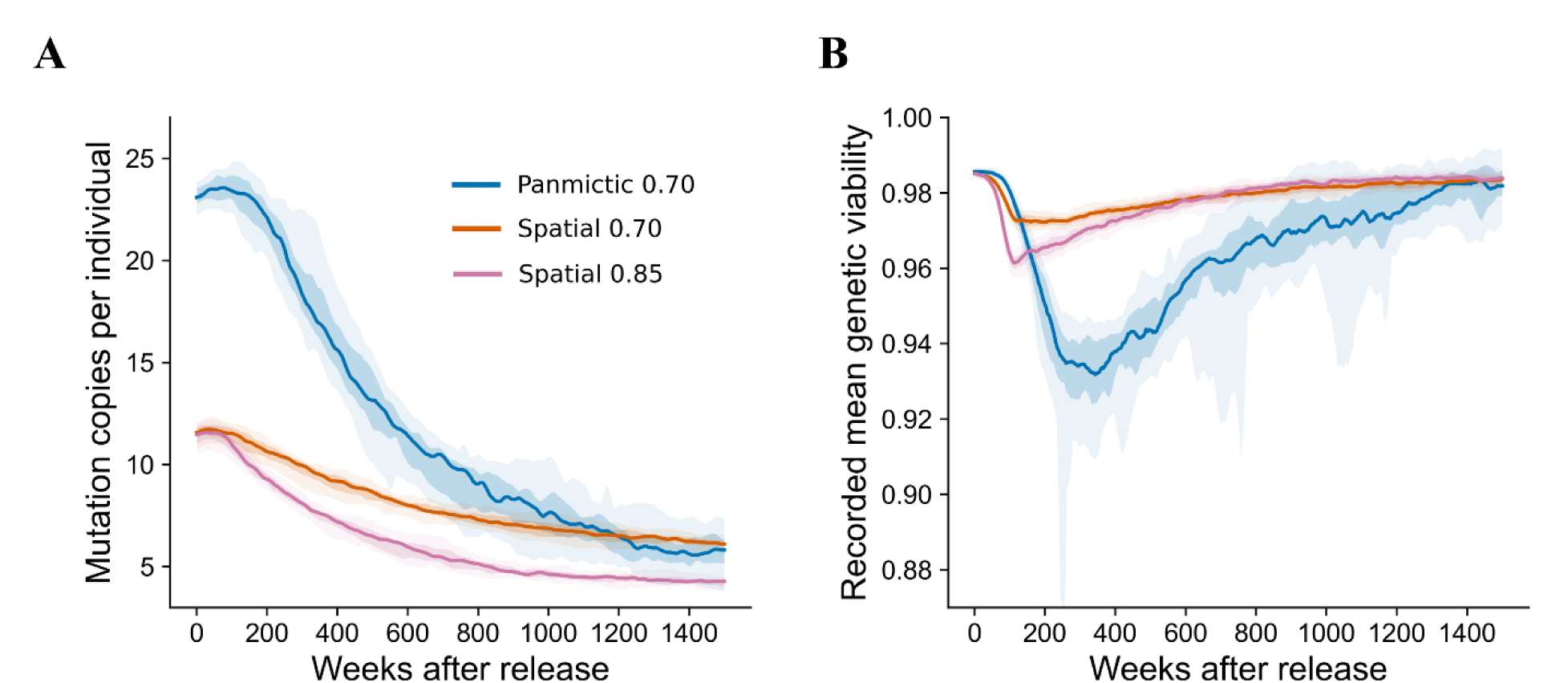
Recessive deleterious load change after drive release. (**A**) Mutation number per in-dividual and (**B**) population mean genetic viability change with time after drive release in the long-term persistence simulations of three settings. The number in the legend represents drive conversion efficiency.

## Supplemental Methods

### S1 Standing variation in drive characteristics

The genome-wide association study on drive conversion and embryo resistance allele formation rate in the *Drosophila* Genetic Reference Panel (DGRP) lines has shown that though these two characteristics both have a genetic basis, there is no common genetic loci of large effect (Champer et al., 2019). Although this result does not establish the genetic architecture of these traits, it is consistent with the possibility that variation is influenced by multiple loci with relatively small individual effects. Such cumulative contributions of small-effect variants have been observed for other quantitative traits (Hua and Springer, 2018). These findings motivated us to consider a polygenic model in which multiple modifier loci contribute additively to standing variation in drive-related traits. Thus, we set *m* loci that have mutations and would affect drive or dispersal characteristics, which are randomly distributed in the genome. Referring to the exponential distribution of effect sizes of tested quantitative traits in yeast (Hua and Springer, 2018), we set the effect size of mutations that affect drive conversion, embryo resistance allele formation rate, and female drive heterozygote fitness to follow a Laplace distribution with a mean equal to 0. The probability density function of the effect size of mutations is:

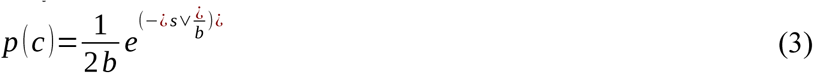

where *s* is the allelic effect and *b* is the Laplace scale parameter. Mutation dosage was additive across allele copies and across loci. For each individual, locus-specific effects were summed with the baseline trait value, and the realized drive conversion probability, embryo resistance probability, and fitness value were each clipped to the interval [0,1].

We assumed all mutations influencing drive conversion, embryo resistance formation rate, and drive heterozygote somatic fitness under no drive release are neutral, so they conform with the neutral theory of molecular evolution and the expected site frequency spectrum following an inverse relationship with allele frequency (Ewens, 2004; Tajima, 1989), which is given as:

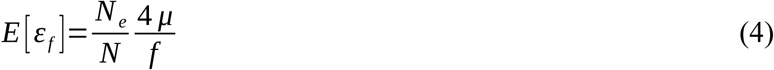

where *εf* represents the mutation number when the mutation frequency is *f*, *Ne* represents the effective population size, *N* represents real population size, and *μ* represents the single nucleotide spontaneous mutation rate. The polymorphism level in *Anopheles stephensi*, *Anopheles coluzzii*, and *Drosophila* have been estimated(Keightley et al., 2014; Rashid et al., 2022), so we set *μ* as 1.36 × 10^-9^. Considering the drive action time is relatively short for the new mutation accumulation, we ignore the occurrence of new mutations that may affect these drive-related properties.

Final realized genetic variation of drive conversion and embryo resistance allele formation rate for all tests in this study were lower than or equal to tested experimental genetic variation in DGRP lines (Champer et al., 2019). This was conservative, especially considering that the genomic background in the DGRP study was only 50% from the DGRP lines and 50% from a laboratory strain background. The initial conditions were controlled such that the discrepancy between the set value and the realized average drive characteristic was maintained below 2%. This controlled for potential outcome bias due to variations in the initial population.

### S2 Mutations affecting dispersal distance

The genetic architecture generation method is the same as mutations affecting drive characteristics. Heritable variation was introduced at initialization using 100 functional mutation loci, with effects drawn from a zero-centered Laplace distribution with scale parameter *b*=0.05. No new dispersal-modifying mutations arose during subsequent simulation; changes in genetic variation therefore reflected the transmission, selection, and loss of standing variants. Independent replicates were initialized by resampling modifier positions, effects, and starting frequencies, together with the initial population. The same genetic-background generation procedure was used across replicates, while the realized genetic backgrounds differed. The modifier number and effect scale were fixed throughout the study. These values defined a specified polygenic scenario and were not treated as empirical estimates of genetic variation in natural mosquito populations. Accordingly, inference was restricted to this initialization procedure without assuming robustness to alternative genetic architectures or levels of standing variation. Female dispersal was specifically used for selecting mates and placing offspring (male dispersal was only used for male movement between time steps).

Under this parameter set, we first evaluated dispersal-dependent fitness costs in populations without a gene drive. We found that populations evolved toward greater dispersal, which most likely slightly increased the chance that they could find a mate or perhaps helped ensure more even offspring distribution with less competition. Thus, we decided to implement a small fitness cost for higher dispersal, representing the energetic cost of movement, aiming to prevent substantial evolution in average dispersal in non-drive control simulations. The direct fitness multiplier was *1-k*d*, where *d* denotes an individual’s dispersal-kernel parameter, and *k* controls the cost strength. This multiplier acted to reduce male mating success rate and adult fecundity from both sexes. Candidate cost strengths were compared using shared initial populations. The test runs comprised a fixed 100-week transition period followed by 1,500 weeks of observation, with 100 replicates per parameter set. The population distribution at the end of the transition period provided the reference for subsequent changes. Control performance was evaluated from trajectories of dispersal mean, standard deviation and distribution tails, population abundance, fertile female abundance, heterozygosity and segregating-locus retention. Following preliminary comparisons, *k*=1.0 was selected for the focal scenario, prioritizing retention of dispersal variation over closer maintenance of the population mean alone. The mean dispersal increased about 1.7% at the endpoint. The average within-population dispersal standard deviation is reduced by 16.89%. Endpoint expected heterozygosity averaged 90.72% of its baseline value. This maintained population abundance and showed a small average displacement in mean dispersal, but exhibited narrowing of the dispersal distribution and minor loss of standing genetic variation. We therefore treated these simulations as an evolving no-drive reference rather than evidence of an invariant trait distribution or genetic equilibrium.

Drive release in dispersal simulations occurred at week 101. Each evolving-dispersal replicate had two matched comparators. A no-drive reference retained heritable dispersal variation and was followed over the corresponding 1,500-week observation period to quantify background dispersal evolution. A fixed-dispersal control received the same drive treatment, but all individuals had a common dispersal parameter equal to the mean across all individuals in the corresponding variable-dispersal population at week 100. The fixed-dispersal control was used to compare drive outcomes. Corresponding control simulations were matched by simulation origin seed.

### S3 Recessive deleterious mutation

#### Mutation input and juvenile viability

All genetic elements in the coarse-grained autosomal genome except the drive locus were potential deleterious-mutation sites and were initially wild type. Each site mutated with probability 2.0 × 10^−6^ per haploid transmission, giving a nominal haploid genomic mutation input of 0.02 per haploid transmission. This value lies within the low-rate, moderate-effect class inferred from *Drosophila melanogaster* mutation-accumulation experiments (García-Dorado et al., 1999; García-Dorado et al., 1998). Because no single experiment jointly estimates the mutation rate, distribution of homozygous effects, dominance, and genomic target used here, we treated these quantities as a literature-informed composite parameterization rather than as a species-specific estimate. For each new mutation, we drew a negative selection coefficient *s* from a reflected Gamma distribution with shape α = 3.35 and rate β = 16.75 (mean α/β = 0.2). The shape parameter was estimated from egg-to-adult viability in *Drosophila melanogaster* mutation-accumulation lines (García-Dorado et al., 2007; García-Dorado et al., 1998), and effects near 0.2 account for a substantial fraction of the mutational decline in fitness components in these experiments (García-Dorado et al., 1999). Values of the selection coefficient were capped at −1 when viability was evaluated, making these rare draws pre-adult lethal. Under the specified Gamma distribution, their probability was approximately 1.63 × 10^-5^.

We focused on completely recessive mutations (h = 0) to isolate the demographic consequences of deleterious variation masked in heterozygotes. Multilocus viability was calculated multiplicatively across expressed homozygous effects (Drake et al., 1998). Thus, for individual *i*,

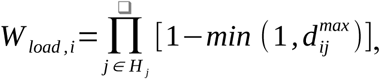

where *H_i_* is the set of expressed positions, and *d_ij_^max^* is the largest decrement carried at position *j* across either homologous chromosome. This position-based rule included the ability to have compound heterozygotes carrying distinct deleterious mutations at the same genomic locus. Mutations confined to one homolog at a position had no direct effect. This independent multiplicative model was used as the null genome-wide fitness model in the absence of locus-specific epistasis (Kondrashov, 1998).

This genetic viability component was applied once to age-0 offspring multiplicatively with the model’s density-dependent juvenile survival factor. Density-dependent competition and survival were evaluated before genotype-based survival was realized, so genetic mortality did not directly reduce larval competition. This represented an aggregate pre-adult viability cost, without separately modelling effects during egg development, hatching, larval development, and pupation. Experimental evidence of inbreeding depression in *Aedes aegypti* supports the biological relevance of juvenile viability effects in mosquitoes (Ross et al., 2019).

#### Virtual mating assay

We assessed potential inbreeding depression for viability by comparing offspring from virtual full-sib matings with offspring from virtual matings between individuals from different families. This comparison measured the proportional viability reduction exposed by the imposed mating pattern, given the population’s current genetic composition. All offspring were virtual and were not added to the simulated population. We performed this assay in each load-bearing population at every reported weekly time step, using 1,000 Monte Carlo pedigrees. For each pedigree, we sampled two distinct females and two distinct males from reproductive adults. One female-male pair produced two virtual full siblings; the second pair produced a reference offspring from a separate family. Virtual meiosis followed the simulation’s recombination map. A gamete from one focal sibling was combined either with a gamete from its full sibling or with a gamete from the reference offspring. The same focal gamete was used in both comparisons. We evaluated the resulting virtual grandchildren using the same *W load* function before selection and without adding them to the simulated population. The assay isolated the deleterious genetic viability component and did not include density-dependent survival or drive-associated fitness effects.

We calculated virtual haploid inbreeding load as

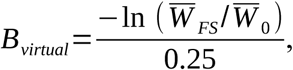

where *W̅_FS_* and *W̅_0_* are the arithmetic mean viabilities of full-sib and random-reference offspring across virtual pedigrees. This is the logarithm of the ratio of mean viabilities, rather than a mean of pedigree-specific log ratios. The denominator is the conventional full-sib offspring inbreeding coefficient under unrelated, non-inbred founders (Szulkin et al., 2007). The logarithmic viability ratio was expressed on the nominal lethal-equivalent scale using the classical relationship between inbreeding coefficient and viability reduction (Duthie et al., 2016; Nietlisbach et al., 2019). This scaling did not correct for relatedness already present among sampled founders. Founders were distinct but were sampled from the current population without excluding existing relatedness. We therefore interpreted *Bvirtual* as an assay-based measure on the nominal haploid lethal-equivalent scale, rather than a pedigree-corrected estimate of realized population inbreeding. It quantifies the additional proportional viability reduction exposed by the imposed pedigree. Larger values indicate greater potential for inbreeding depression.

#### Standing variation and pre-release stability

Populations were evolved for 6,000 weeks before drive release to establish standing mutation-selection-drift variation and were followed until week 7500. New deleterious mutations continued to arise after release, allowing post-release dynamics to include both standing and newly arising variation. Convergence in the final simulations was assessed from 250-week block means over the final 1,000 pre-release weeks for virtual inbreeding load, deleterious copies per individual, and population abundance. For each metric, we calculated the absolute fitted change projected across the 1,000-week interval and the absolute difference between the final two block means. Both quantities were expressed relative to the mean in the final block. A metric was considered operationally stable when both relative changes were below 5%.

### S4 Statistical procedures

#### Analysis units and endpoint definitions

Independent simulations were the units of analysis. Functional and neutral drive-characteristic simulations were unpaired, as were simulations with and without recessive deleterious variation. Dispersal simulations sharing an origin seed were paired. Repeated weekly observations were summarized within each simulation before comparisons, and outcome probabilities used all simulations with classifiable outcomes.

For drive-characteristic analyses, the final trait value was the adult-abundance-weighted average of weekly population mean traits over the final 20 weeks. Final relative population size was the mean abundance over the final 20 weeks divided by carrying capacity. Group comparisons included independent simulations with a defined endpoint rather than being restricted automatically to long-term-persistence outcomes.

For recessive-load analyses, minimum relative population size was the minimum abundance over the complete post-release trajectory divided by the mean abundance over the final 20 pre-release time steps. Complete elimination contributed a minimum of zero. Final relative population size was the mean abundance over the final 20 weeks divided by the mean over the final 20 pre-release weeks. Each simulation was normalized to its own pre-release baseline.

#### Independent continuous and outcome comparisons

We compared continuous endpoints within each regime using two-sided independent two-sample permutation tests. The statistic and effect estimate were the difference in arithmetic means. Under the exchangeability null, the two groups followed the same distribution. We pooled their endpoints and reassigned treatment labels 10,000 times while retaining the original group sizes. The two-sided P value was the fraction of permuted absolute mean differences at least as large as the observed absolute difference.

P values were Holm-adjusted across eligible regimes separately for each continuous endpoint within the relevant modifier or load analysis. Comparisons with fewer than two observations in either group were descriptive. Group means and percentile-bootstrap 95% confidence intervals were obtained from 10,000 within-group resamples of independent simulations.

Selected unpaired outcome contrasts used two-sided Fisher’s exact tests, coding the focal outcome against all other outcomes. These exploratory contrasts were selected after inspection of outcome distributions and were reported without multiplicity adjustment. The recessive-load elimination contrast used the same test. Outcome probabilities were reported with Wilson 95% confidence intervals.

#### Associations across spatial settings

We analyzed embryo resistance and female drive-heterozygote fitness separately across seven spatial settings. For each setting, the baseline spatial metric was the neutral-control mean NNI over the final 20 weeks, and the response was the corresponding variable-treatment mean final trait. Setting-level means and 95% confidence intervals were estimated from 10,000 bootstrap resamples of independent simulations. Two-sided exact Spearman *P* values were calculated by enumerating the permutation distribution of the seven setting-level pairings. The setting, not the individual simulation, was the correlation unit. *P* values were unadjusted, and associations were not interpreted as demonstrating causality.

Figure S1 separately displays final-20-week NNI distributions for selected modifier settings. These distributions describe spatial outcomes within treatments and should be distinguished from the setting-level correlation unit and window defined above.

#### Standing variation and genetic architecture

Realized pre-release variation was the mean adult trait standard deviation over the 10 time steps immediately before drive release. Conversion standard deviation was calculated across all adults. For embryo resistance and female drive-heterozygote fitness, we calculated each adult male’s potential trait from his genotype as if expressed in a female and included these values with adult female values. This measured genetic variation carried by the adult population without treating male values as observed female phenotypes.

We compared a fixed-m path (m = 50, varying b) with a fixed-b path (the trait-specific minimum b, varying m). Each original setting contained 100 simulations. The shared m = 50, b = b_min_ setting was excluded from both comparison groups. Models were fitted only within empirical common support, the intersection of the two observed realized-standard-deviation ranges.

We centered realized standard deviation at the common-support median and scaled it in 0.01-unit increments. P_i_ indicated path, with 0 for fixed m and 1 for fixed b. The final trait endpoint Y_i_ was modeled as:

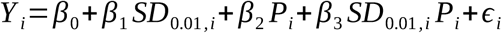

Coefficients were estimated by ordinary least squares. Standard errors, 95% confidence intervals, and joint Wald tests used the HC3 heteroskedasticity-robust covariance matrix. The realized-variation test evaluated β₁ = β₃ = 0, equivalent to zero slopes along both paths. The architecture test evaluated β₂ = β₃ = 0, equivalent to no adjusted endpoint offset and no slope difference between paths. β₃ estimated the path-specific slope difference, and β₂ estimated the adjusted difference at the reference standard deviation.

The two primary test families were Holm-adjusted separately across the three traits. Component tests of slope differences and reference-standard-deviation offsets were reported with unadjusted P values.

#### Matched dispersal comparisons

Evolving- and fixed-dispersal outcomes were matched by origin seed. At drive conversion efficiencies of 0.95 and 0.99, elimination and drive loss were tested separately against all other outcomes using two-sided exact McNemar tests. Inference used the two directional discordant-pair counts. Comparisons with no discordant pairs were assigned *P* = 1. The four outcome-by-setting comparisons formed one Holm family. Outcomes at 0.75 and 0.85 were descriptive.

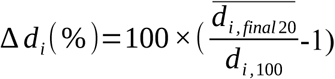

Cumulative adult dispersal change was calculated by comparing mean adult dispersal over the final 20 weeks with its week-100 baseline. Evolving-drive simulations were divided by their final outcomes into long-term-persistence and non-long-term-persistence groups. Within each setting and group, we compared change with origin-seed-matched no-drive references using a two-sided paired t-test. The null was a zero mean within-pair difference. Effects were reported as arithmetic mean paired differences in percentage.

Empty outcome groups were not tested. P values were Holm-adjusted across all eligible setting-by-outcome-group comparisons, separately from the outcome-test family. Percentile-bootstrap 95% confidence intervals used 10,000 origin-seed resamples, retaining complete pairs. Because outcome groups were defined after the simulations, their contrasts were interpreted with regard to outcome selection and differing termination times.

#### Trajectories and reporting

Spatial NNI trajectories were summarized by across-simulation medians and interquartile ranges. Virtual inbreeding-load trajectories in Figure 6C were summarized among load-bearing simulations ending in long-term persistence using medians and central 50% and 90% intervals. These bands described variation among simulations, not confidence intervals for the median. No formal trajectory tests were performed. In Figure S3, dark and light bands denote the interquartile range and 5th–95th percentiles, respectively, and all panels describe the same no-drive references. Statistical significance was assessed at P < 0.05, using adjusted P values for the specified test families. Exact sample sizes, effect estimates, and raw and adjusted P values were reported in Supplementary Tables and Source Data. Matched outcome analyses additionally reported the two directional discordant-pair counts.

